# TDP-43 oligomerization and RNA binding are codependent but their loss elicits distinct pathologies

**DOI:** 10.1101/2022.05.23.493029

**Authors:** Manuela Pérez-Berlanga, Vera I. Wiersma, Aurélie Zbinden, Laura De Vos, Ulrich Wagner, Chiara Foglieni, Izaskun Mallona, Katharina M. Betz, Antoine Cléry, Julien Weber, Zhongning Guo, Ruben Rigort, Pierre de Rossi, Ruchi Manglunia, Elena Tantardini, Sonu Sahadevan, Oliver Stach, Marian Hruska-Plochan, Frederic H.-T. Allain, Paolo Paganetti, Magdalini Polymenidou

## Abstract

Aggregation of the RNA-binding protein TDP-43 is the main common neuropathological feature of TDP-43 proteinopathies. In physiological conditions, TDP-43 is predominantly nuclear and contained in biomolecular condensates formed via liquid-liquid phase separation (LLPS). However, in disease, TDP-43 is depleted from these compartments and forms cytoplasmic or, sometimes, intranuclear inclusions. How TDP-43 transitions from physiological to pathological states remains poorly understood. Here, we show that self-oligomerization and RNA binding cooperatively govern TDP-43 stability, functionality, LLPS and cellular localization. Importantly, our data reveal that TDP-43 oligomerization is connected to, and conformationally modulated by, RNA binding. Mimicking the impaired proteasomal activity observed in patients, we found that TDP-43 forms nuclear aggregates via LLPS and cytoplasmic aggregates via aggresome formation. The favored aggregation pathway depended on the TDP-43 state –monomeric/oligomeric, RNA-bound/-unbound– and the subcellular environment –nucleus/cytoplasm. Our work unravels the origins of heterogeneous pathological species occurring in TDP-43 proteinopathies.

## Introduction

Amyotrophic lateral sclerosis (ALS) and frontotemporal lobar degeneration (FTLD) are two seemingly different, devastating adult-onset neurodegenerative diseases that exhibit a significant genetic, clinical and pathological overlap (*1*). The vast majority of ALS patients and up to half of FTLD cases are characterized by the accumulation of aggregated TAR DNA-binding protein 43 (TDP-43) in affected neurons (*1*–*3*). Importantly, TDP-43 pathology is not an exclusive hallmark of ALS and FTLD, but is also the main pathological feature in limbic-predominant age-related TDP-43 encephalopathy (LATE) (*4*) and a concomitant pathology in a subset of patients with other neurodegenerative diseases including Alzheimer’s, Parkinson’s and Huntington’s disease (*5*). However, TDP-43 aggregates associated with different clinical subtypes present distinct subcellular localization, namely cytoplasmic, intranuclear or axonal (*6, 7*), as well as morphological and biochemical properties (*8*–*10*), indicating a distinct molecular origin of these diverse pathological species.

TDP-43 is a ubiquitously expressed (*11*) and predominantly nuclear (*12*) nucleic acid-binding protein (*11, 13*) composed of an N-terminal domain (NTD, amino acids 1-80) involved in self-oligomerization (*14, 15*), two tandem RNA-recognition motifs (RRMs, amino acids 106-259) (*13, 16*) and an unstructured low complexity region (LCR, amino acids 260-414). The latter contains a transient α-helix (amino acids 321-340) (*17*) that associates with interaction partners (*18*) and was recently shown to coincide with the aggregation core of pathological cytoplasmic TDP-43 in FTLD brains (*19*). Under physiological conditions in the nucleus, TDP-43 mainly binds UG-rich intronic sites on pre-mRNA to regulate alternative splicing (*20, 21*) and undergoes liquid-liquid phase separation (LLPS) (*22*) to form dynamic nuclear droplets (*23*–*26*), which were suggested to localize to specific subnuclear membraneless compartments (*26, 27*). Albeit its predominantly nuclear localization (*12*), TDP-43 shuttles between the nucleus and the cytoplasm (*28*), where it plays roles in mRNA stability, transport and translation, miRNA processing, mitochondrial and synaptic function and stress responses (*1*). In particular, TDP-43 was shown to incorporate into and modulate the dynamics of stress granules (SGs) upon exposure to different temperature, osmotic, oxidative and chemical stressors (*29, 30*).

RRM-mediated RNA binding is essential for TDP-43 to perform its physiological functions in RNA metabolism (*31*). In addition, RNA binding precludes TDP-43 passive leakage out of the nucleus (*28, 32*) and modulates its LLPS behavior (*33, 34*). In contrast, little is known about the importance of TDP-43 NTD-driven self-oligomerization in physiology. Previous data have shown that nuclear TDP-43 oligomerization is required for alternative splicing of at least a subset of its known RNA targets (*15, 35*–*38*). However, the role of TDP-43 oligomerization in its physiological properties, including its subcellular localization, stability, LLPS behavior and cytoplasmic functions, remains poorly understood. Also, whether -and if so, how-TDP-43 RNA binding and oligomerization impact each other in cells is unknown.

The overexpression of TDP-43 in cellular and animal models results in its aggregation, a phenomenon that has been extensively explored in recent years to recapitulate the main neuropathological hallmark of ALS/FTLD (*39*). However, TDP-43 is a tightly autoregulated protein (*20, 40*) and overexpression can distort its subcellular (*32*) and subnuclear (*24, 27*) localization, and potentially its functions. We therefore aimed to study the physiological role of TDP-43 oligomerization and its interplay with RNA binding at near-physiological protein levels, and to subsequently compare the pathways triggered by their respective impairment. Using human neural cultures and single-copy expression systems in human cell lines, we show that NTD-driven TDP-43 oligomerization and RRMs-mediated RNA binding are intertwined and required to maintain the half-life, functionality and localization of TDP-43. Upon failure of the ubiquitin-proteasome system (UPS), monomerization and impaired RNA binding triggered TDP-43 aggregation via distinct pathways in the cytoplasm and nucleus. Our results underscore the relevance of loss of oligomerization and RNA binding in the initiation of diverse TDP-43 pathologies and unravel the origins of heterogeneous pathological species occurring in human disease.

## Results

### Oligomerization and RNA-binding cooperatively stabilize the half-life of TDP-43

To systematically compare the properties of oligomeric, monomeric and RNA binding-deficient TDP-43, we introduced a single copy of an N-terminally green fluorescent protein (GFP)-tagged TDP-43 coding sequence under the control of a doxycycline-inducible promoter into HEK293 cells using the Flp-In T-REx technology (**Figure S1A**) (*41*). This construct harbored previously described point mutations that disrupt TDP-43 oligomerization (termed 6M) (*15*), RNA binding through the RRMs (five F>A mutations in the RRMs, referred to as RRMm) (*13, 16*) or both 6M&RRMm (**Figure 1A**). The resulting four isogenic cell lines (WT, 6M, RRMm and 6M&RRMm) expressed equal levels of the exogenous GFP-TDP-43 RNA (**Figure S1B**) and both RNA and protein were detectable only upon addition of doxycycline (**Figures 1B and S1C-D**). Wild type (WT) GFP-TDP-43 protein levels displayed a mere 4-fold increase compared to endogenous TDP-43, as determined by immunoblot analysis (**Figure S1D**). However, despite equal RNA levels (**Figure S1B**), protein levels of the GFP-TDP-43 mutants were noticeably lower than their WT counterpart (**Figures 1B-C and S1E-G**). A protein turnover analysis using the translation inhibitor cycloheximide (CHX) showed that the half-life of the RNA-binding TDP-43 mutant was reduced by >8 hours compared to WT GFP-TDP-43 (**Figure 1D-E**), consistent with previous findings (*42*). Interestingly, oligomerization deficiency (6M) and loss of RNA binding had a similar effect on the half-life of TDP-43. Furthermore, the combined variant (6M&RRMm) presented a cumulative effect (**Figure 1D-E**). Since point mutations can affect protein folding and thereby selectively target proteins for degradation (*43*), we confirmed that the introduced mutations do not interfere with the folding of TDP-43 using circular dichroism (CD) (**Figure 1F and S1H**) and two-dimensional nuclear magnetic resonance (2D-NMR) spectroscopy (**Figure S1I-J**) (*15, 44*), which revealed that the mutated domains are properly folded. Therefore, our results indicate that loss of oligomerization or RNA-binding ability similarly reduces the half-life of TDP-43. Since incorporation of proteins into multimeric complexes has been reported to correlate with longer half-lives in yeast (*45*) and mouse brain cells (*46*), these observations strengthen the link between TDP-43 functionality and its half-life.

**Figure 1.**
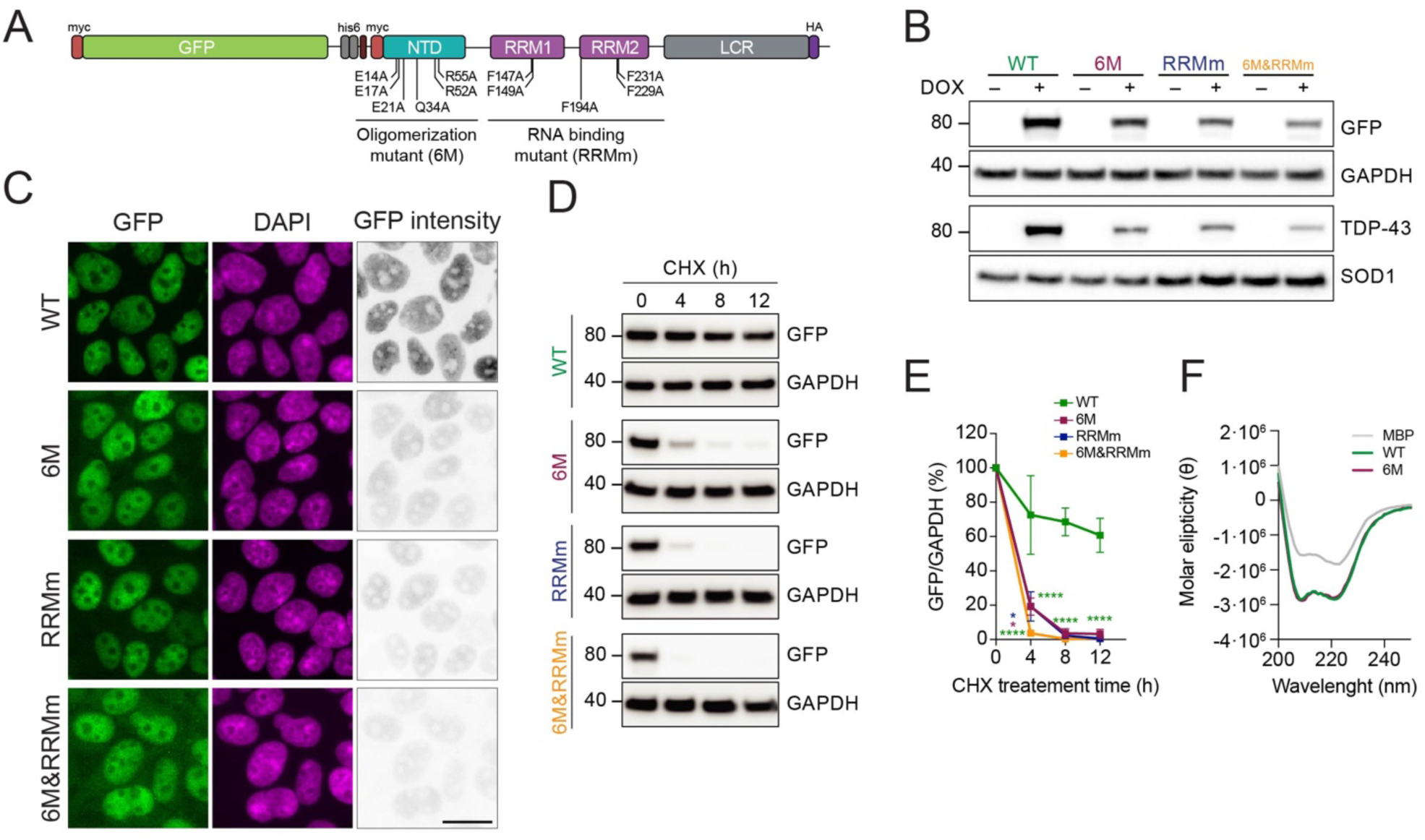
Oligomerization and RNA-binding cooperatively stabilize the half-life of TDP-43. **(A)** Schematic representation depicting the specific mutations used to disrupt oligomerization and/or RNA binding on the GFP-TDP-43 variants, used to develop the inducible, isogenic cell lines. **(B)** Western blot analysis of the generated isogenic cell lines described in A after inducing GFP-TDP-43 expression for 48 h showing the tightness of the doxycycline (DOX)-modulated expression system. Note also the different protein levels of the expressed variants. **(C)** Representative images of widefield fluorescence microscopy of the isogenic HEK293 cell lines depicted in B. GFP brightness is adjusted in each condition for optimal visualization of GFP-TDP-43 localization. Original intensity values are represented in the right column using grayscale. Scale bar: 20 µm. **(D)** GFP-TDP-43 expression was induced with DOX for 24 h before cycloheximide (CHX) treatment for the indicated times and western blot analysis. **(E)** Quantification of the GFP signal from D. N=3 independent experiments. Two-way ANOVA with Tukey’s multiple comparisons post hoc test. **(F)** Average far-UV CD spectra of purified TDP-43-MBP variants from N=3 independent experiments. **** p<0.0001. Graph bars represent mean ± SD.

### TDP-43 oligomerization and RNA-binding preserve its nuclear localization

Due to an active nuclear import and its ability to passively diffuse out of the nucleus, TDP-43 is a nucleocytoplasmic shuttling protein (*32, 47*). RNA binding retains TDP-43 in the nucleus by forming bigger macromolecular complexes that slow down its diffusion (*28, 32*). We therefore wondered whether oligomerization is a greater driver of physiological TDP-43 nuclear localization. To address this question, we first measured the mean fluorescence intensity of GFP-TDP-43 in the nucleus and the cytoplasm for all four GFP-TDP-43 variants. Using DNA and G3BP as nuclear and cytoplasmic markers, respectively (**Figure S2A**), we observed that monomeric TDP-43 (6M and 6M&RRMm) showed a significantly increased cytoplasmic localization compared to WT GFP-TDP-43 (**Figure 2A and S2A**). Mislocalization was exacerbated in combination with a loss of RNA binding, suggesting an independent, additive contribution of both protein-protein and protein-RNA interactions to the nuclear localization of TDP-43. Similar observations were obtained in human neurons transduced with HA-tagged versions of the TDP-43 variants (**Figures 2B-C**).

**Figure 2.**
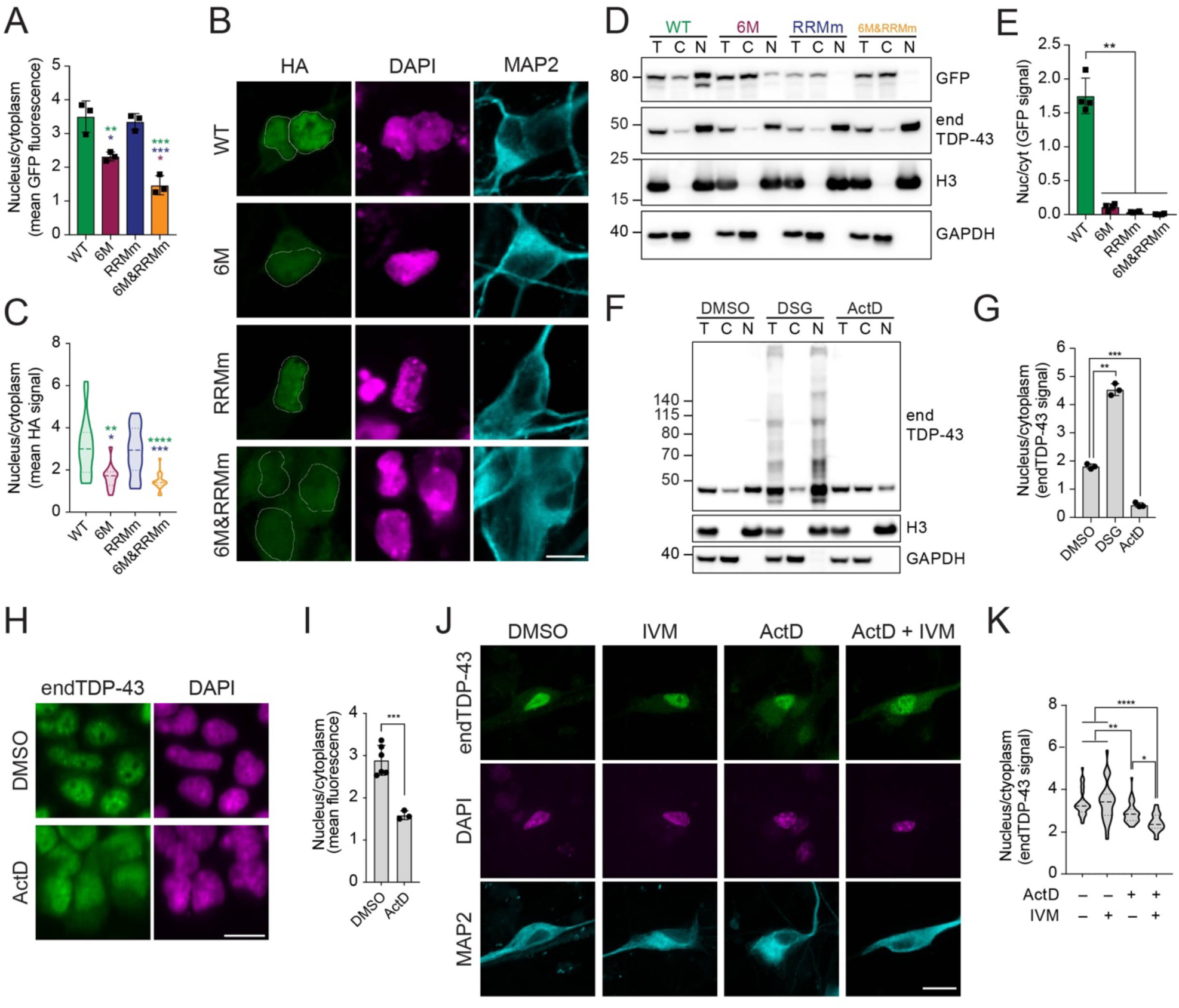
TDP-43 oligomerization and RNA-binding preserve its nuclear localization. **(A)** Quantification of nucleocytoplasmic levels of GFP-TDP-43 in the immunocytochemistry images shown in Figure 1C. N=3 independent experiments. One-way ANOVA with Tukey’s multiple comparisons post hoc test. **(B)** Representative image of confocal fluorescence imaging of human neurons transduced with TDP-43-HA variants. Scale bar: 10 µm. **(C**) Quantification of nucleocytoplasmic levels of TDP-43-HA in the immunocytochemistry images shown in B. N=14-20 cells. Kruskal-Wallis test with Dunn’s multiple comparisons post hoc test. **(D)** GFP-TDP-43 expression was induced with doxycycline (DOX) for 4 h before nucleocytoplasmic fractionation and subsequent analysis of GFP-TDP-43 and endogenous TDP-43 (endDP-43) by western blot. T: total lysate, C: cytoplasmic fraction, N: nuclear fraction. **(E)** Quantification of the GFP signal from Figure 2D. Repeated measures one-way ANOVA with Greenhouse-Geisser correction and Tukey’s multiple comparisons post hoc test. cyt: cytoplasm, nuc: nucleus. **(F)** Representative images of widefield fluorescence microscopy of HEK293 cells treated with ActD for 4 h. Scale bar: 20 µm. **(G)** Quantification of F. N=3 independent experiments. Unpaired two-tailed *t*-test. **(H)** HEK293 cells were treated with ActD to inhibit transcription or subjected to protein-protein cross-link with DSG followed by nucleocytoplasmic fractionation and western blot analysis. T: total lysate, C: cytoplasmic fraction, N: nuclear fraction. **(I)** Quantification of endTDP-43 signal shown in H. N=3 independent experiments. Repeated measures one-way ANOVA with Greenhouse-Geisser correction and Dunnett’s multiple comparisons post hoc test. **(J)** Representative images of confocal fluorescence microscopy of human neural cultures treated with actinomycin D (ActD) and ivermectin (IVM). Scale bar: 20 µm. **(K)** Quantification of nucleocytoplasmic levels of endTDP-43 in the immunocytochemistry images shown in J. Kruskal-Wallis test with Dunn’s multiple comparisons post hoc test. N=23-48 fields corresponding to a total of 351-569 neurons per condition. * p<0.05, ** p<0.01, *** p<0.001, **** p<0.0001. Graph bars represent mean ± SD. Violin plots show mean and quartiles.

We subsequently sought to confirm these results by nucleocytoplasmic fractionation. In line with the immunocytochemistry results (**Figure 1C and 2A-C**), upon mild lysis and nuclei isolation by centrifugation, WT GFP-TDP-43 was mostly retained in the nuclear fraction (**Figure 2D-E**), with a small cytoplasmic pool corresponding to the subset of TDP-43 performing functions in this compartment (*1*). In contrast, oligomerization-deficient mutations (6M and 6M&RRMm) consistently shifted the majority of GFP-TDP-43 to the cytoplasmic fraction (**Figures 2D-E**), to a larger extent than observed by immunocytochemistry (**Figure 2A and 2C**). Importantly, the endogenous protein in the same samples remained predominantly nuclear (**Figure S2B**). This contrast was even more pronounced in the case of RRMm GFP-TDP-43, which fully shifted its localization to the cytoplasm upon fractionation (**Figures 2D-E**) as opposed to its nuclear localisation by immunocytochemistry (**Figure 2A and 2C**). We therefore wondered whether monomeric and RNA binding-deficient GFP-TDP-43 exhibit an increased passive diffusion rate and diffuse out of the nucleus during the fractionation procedure, when active nuclear import is absent. Indeed, stabilization of TDP-43 oligomers by protein-protein cross-linking with disuccinimidyl glutarate (DSG) (*15*) before nucleocytoplasmic fractionation increased the retention of endogenous TDP-43 in the nucleus (**Figure 2F-G**). Conversely, when we pretreated the cells with actinomycin D (ActD) to block transcription, the localization of endogenous TDP-43 shifted to the cytoplasm (**Figure 2F-G and S2C-D**), as previously reported (*28, 32*). Notably, this efflux of TDP-43 upon a decrease in nuclear RNA levels and subsequent sample fractionation (**Figure 2F-G**) was more pronounced than observed by immunocytochemistry (**Figure 2H-I**). These observations suggest that active nuclear import compensates for the abundant passive TDP-43 egress from the nucleus in the absence of oligomerization or RNA binding. Similar results were observed in human neurons, where the combined treatment of ActD and ivermectin, a nuclear import inhibitor, increased the cytoplasmic shift of endogenous TDP-43 as compared to treatment with ActD alone (**Figure 2J-K**). Collectively, these observations show that RNA binding and protein-protein interactions, especially its self-oligomerization, involve TDP-43 in larger macromolecular complexes that are retained in the nucleus.

### Oligomerization is required for physiological phase separation of TDP-43 in the nucleus

TDP-43 has been shown to undergo LLPS (*22*), a phenomenon visible in the nucleus as small droplets that fuse and split at endogenous protein concentrations (*23*–*26*). Such endogenous LLPS-driven droplets were also observed in our model (**Figure 3A**) as confirmed by treatment with 1,6-hexanediol (1,6-HD), an LLPS-suppressing alcohol (*48*). 1,6-HD decreased both the number and size of endogenous TDP-43 droplets (**Figure 3A-B and S3A**). Although high concentrations of the LCR of TDP-43 are sufficient for phase separation *in vitro* (*17, 49*), additional interactions must take place for full-length TDP-43 to undergo physiological LLPS at far lower concentrations (*24, 38*). Recent evidence points towards self-interaction through the NTD as another driver of TDP-43 LLPS *in vitro* (*37, 38*) and in human cells (*34*). Indeed, at a reported physiological concentration of 10 µM (*25, 50*) and using dextran as a crowding agent (*25, 51*) purified maltose binding protein (MBP)- tagged full-length TDP-43 phase separated into droplets, which also dissolved in the presence of 1,6-HD (**Figure 3C-D**). In contrast, oligomerization-deficient TDP-43 (6M) did not form droplets under the same conditions, suggesting that NTD interactions are essential for TDP-43 LLPS (**Figure 3C-D**).

**Figure 3.**
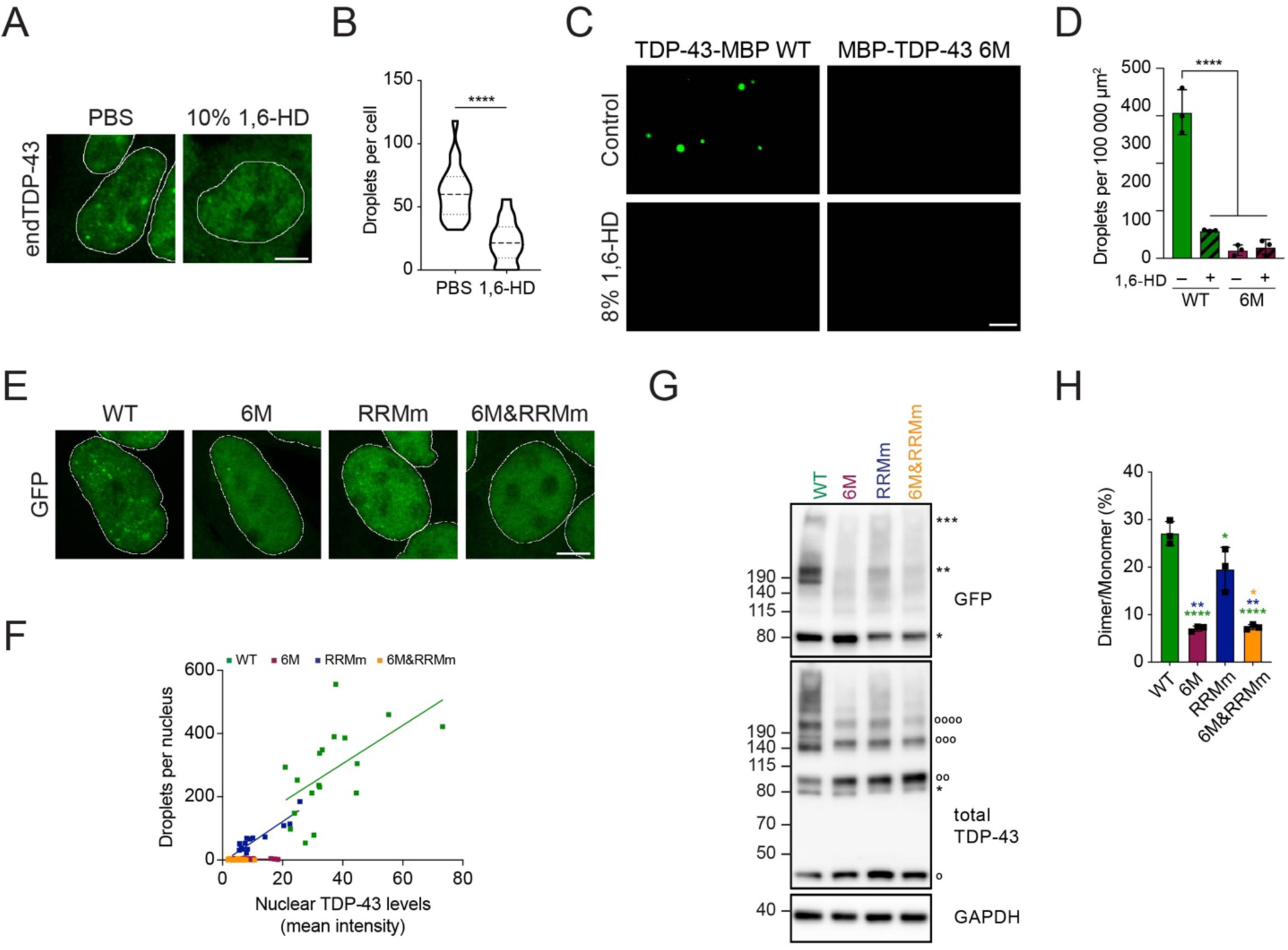
Oligomerization is required for physiological LLPS of TDP-43 in the nucleus. **(A)** Representative maximum intensity Z-projections from confocal fluorescence imaging (thickness of ∼10 µm, in steps of 0.21 µm) of WT HEK293 cells after mock or 1,6-HD treatment for 15 min stained for endogenous TDP-43 (endTDP-43). Scale bar: 5 µm. **(B)** Quantification of three-dimensional (3D) reconstructions from the Z-stack confocal microscopy images shown in A depicting the number of nuclear droplets per cell in the conditions described in A. N=14-23 cells. Unpaired two-tailed *t*-test. **(C)** Fluorescence microscopy images of 10 μM purified MBP-tagged full-length TDP-43 and its oligomerization-deficient counterpart showing different abilities to undergo LLPS and its disruption by 1,6-HD treatment for 10 min. Scale bar: 10 μm. **(D)** Quantification of the number of droplets in the conditions shown in C per 100 000 µm^2^ field. N=3 independent experiments. One-way ANOVA with Tukey’s multiple comparisons post hoc test. **(E)** Representative maximum intensity Z-projections (thickness of ∼10 µm, in steps of 0.21 µm) from confocal fluorescence microscopy of the isogenic cell lines expressing GFP-TDP-43 for 48 h with doxycycline (DOX). Scale bar: 5 µm. **(F)** 3D quantification of the number of nuclear droplets per cell shown in E. N=16-22 cells. **(G)** GFP-TDP-43 expression was induced with DOX for 4 h before crosslinking protein-protein interactions with DSG and subsequent analysis by western blot. *, ** and *** indicate GFP-TDP-43 monomers, dimers and trimers, respectively. ^o, oo, ooo^ and ^oooo^ indicate endTDP-43 monomers, dimers, trimers and tetramers. **(H)** Quantification of GFP-TDP-43 dimer/monomer ratio based on the GFP signal from G. N=3 independent experiments. Repeated measures one-way ANOVA with Greenhouse-Geisser correction and Tukey’s multiple comparisons post hoc test. * p<0.05, ** p<0.01, **** p<0.0001. Graph bars represent mean ± SD. Violin plots show mean and quartiles.

These findings were reproduced in our isogenic cell lines. At comparable protein levels, oligomerization disruption virtually suppressed all GFP-TDP-43 droplet formation (**Figures 3E-F**), pointing to an essential role for oligomerization in physiological TDP-43 LLPS in cells. Interestingly, RNA binding-deficient GFP-TDP-43 also formed nuclear droplets (**Figures 3E-F**), albeit smaller in size (**Figure S3B**). Since local protein concentration is a known driver of phase separation (*22*), the number of GFP-TDP-43 droplets for both WT and RRMm was proportional to their nuclear protein levels (**Figure 3F**). Disruption of both oligomerization and RNA binding in the combined GFP-TDP-43 variant drastically reduced the number of nuclear droplets (**Figure 3E-F**), indicating that TDP-43 droplet formation in the absence of RNA binding is mediated through NTD-driven TDP-43 oligomerization. This was supported by biochemical analysis with DSG protein-protein crosslinking which showed protein complexes at the expected size of GFP-TDP-43 dimers for both WT and RRMm, but not for the 6M-containing variants (**Figures 3G-H and S3C-D**). Similar to its WT counterpart, stabilization of these protein complexes via crosslinking retained GFP-TDP-43 RRMm predominantly in the nucleus in our isogenic cell lines despite the lack of RNA binding (**Figure S2C-D**). Overall, our results indicate that NTD-driven oligomerization –and not only LCR interactions– are essential for TDP-43 LLPS in cells, both in the presence and absence of RNA.

### Loss of RNA binding leads to conformationally distinct TDP-43 oligomers

The observed GFP-TDP-43 dimers in the RRMm variant were significantly reduced compared to TDP-43 with retained RNA binding capability (**Figures 3G-H**), suggesting that TDP-43 oligomerization is modulated by RNA binding. Indeed, treatment with ActD decreased the level of endogenous TDP-43 dimers detected by protein-protein cross-linking (**Figure 4A-B**). Concomitant with this reduction in oligomerization, ActD-treated cells also displayed a reduced number of TDP-43 nuclear droplets (**Figure 4C-D**). To determine whether TDP-43 oligomerization is exclusively confined to nuclear droplets, we first assessed the exact subnuclear location of the oligomers. For this purpose, we employed proximity ligation assay (PLA) to visualize TDP-43 dimers with a single monoclonal antibody (mAb) conjugated to two different oligonucleotides. With this approach, only TDP-43 molecules that come to close proximity (maximum 20 nm apart) allow oligonucleotide hybridization and fluorescent signal amplification. Surprisingly, while abundant PLA signal was detected in the nucleus, only a fraction overlapped with its nuclear droplets, suggesting that TDP-43 dimerization is not restricted to nuclear droplets (**Figure 4E**), a result that was confirmed for WT GFP-TDP-43 using a mAb against GFP (**Figure 4F and S4A**).

**Figure 4.**
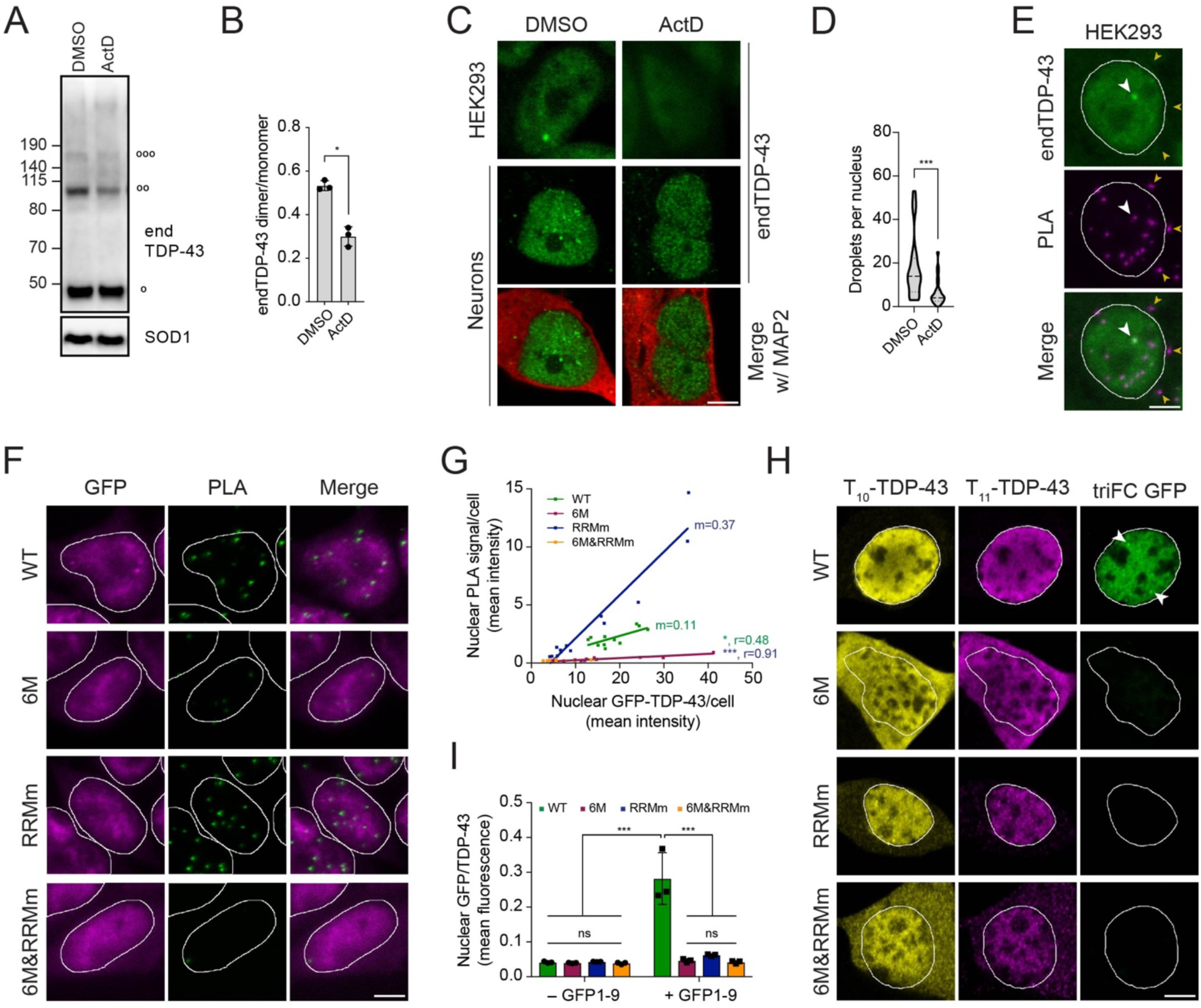
Loss of RNA binding leads to conformationally distinct TDP-43 oligomers. **(A)** HEK293 cells were treated with ActD for 4 h to inhibit transcription before treatment with the protein-protein cross-linker DSG and western blot analysis. ^o, oo^ and ^oooo^ indicate endogenous TDP-43 (endTDP-43) monomers, dimers and trimers, respectively. **(B)** Quantification of the endTDP-43 signal from A. N=3 independent experiments. Paired two-tailed *t*-test. **(C)** Representative image of confocal fluorescence microscopy of HEK293 cells and neurons treated with ActD. Scale bar: 5 µm. **(D)** Single-plane quantification of the number of nuclear droplets per neuron in the conditions described in C. N=25-26 cells. Mann-Whitney U test. **(E)** Proximity ligation assay (PLA) using a monoclonal anti-TDP-43 antibody reveals nuclear and cytoplasmic localization of endTDP-43 dimers in physiological conditions. Big white arrowheads indicate overlapping GFP-TDP-43 droplets and PLA signal. Small yellow arrowheads indicate cytoplasmic PLA signal. Scale bar: 5 µm. **(F)** PLA using a monoclonal anti-GFP antibody reveals the localization of GFP-TDP-43 dimers in the isogenic cell lines upon protein expression with doxycycline (DOX) for 48 h. Note the absence of dimers in the oligomerization-deficient variant (6M). Scale bar: 5 µm. **(G)** Quantification of the nuclear PLA signal shown in G correlated to the protein expression levels of the different TDP-43 variants, measured as the mean GFP fluorescence. N=11-13 cells. **(H)** Tripartite GFP complementation assay using a pair of N-terminally T_10_- and T_11_-tagged TDP-43 constructs co-transfected with GFP_1-9_ in motoneuron-like NSC-34 cells. triFC: trimolecular fluorescence complementation. Scale bar: 5 µm **(I)** Quantification of the GFP fluorescence levels relative to the T_10_/T_11_-TDP-43 expression levels as shown in H. N=3 replicates, with N=6-35 cells per replicate. Two-way ANOVA with Tukey’s multiple comparisons post hoc test. * p<0.05, *** p<0.001. Graph bars represent mean ± SD. Violin plots show mean and quartiles.

In contrast, the oligomerization-deficient variants showed a markedly decreased PLA signal, even in cells with comparable protein levels, confirming that the observed PLA signal depends on oligomerization. In line with our DSG cross-linking results (**Figure 3G-H**), the RNA-binding GFP-TDP-43 mutant displayed a positive PLA signal (**Figure 4F**).

Interestingly, the mean intensity of PLA-positive foci was consistently higher than that of its WT counterpart (**Figure 4F**) at comparable protein levels (**Figure 4G**). These results were confirmed using a mAb targeting a different tag in the GFP-TDP-43 protein (**Figure S4B**). We further validated this finding with an alternative approach using a GFP tripartite fluorescence complementation (triFC) assay that measures physiological dimerization, as we previously showed (*31, 52*) (**Figure S4C**). Co-transfection of T_10_- and T_11_-tagged TDP-43 variants along with a nuclear-targeted GFP_1-9_ in mouse motor neuron-like cells showed positive GFP complementation signal for the WT protein, but none of the other variants (**Figure 4H-I**), despite similar nuclear protein levels (**Figure S4D-E**). This suggests that despite our biochemical (**Figures 3G-H**) and imaging (**Figures 4F-G and S4B**) observations indicating that RNA binding-deficient TDP-43 dimerizes, these dimers do not come in close enough contact and with the correct orientation to reconstitute GFP fluorescence. This contrast to the WT protein supports the notion of a distinct conformation of TDP-43 dimers in the absence of RNA. Of note, transient expression of the monomeric GFP-TDP-43 variants in this cellular model also showed a noticeable cytoplasmic fraction (**Figures 4H and S4F**), comparable to our observations in the isogenic cell lines and in human neurons (**Figure 2B and C**). Overall, detection and quantification of dimeric TDP-43 species by a combination of different imaging and biochemical methodologies supports the view that RNA binding is required for the proper orientation of TDP-43 dimers and likely oligomers.

### TDP-43 partitions in heterogenous nuclear bodies via oligomerization and RNA binding

Since RNA promotes phase separation by providing a scaffold for many phase-separated nuclear bodies (*53, 54*), we sought to determine whether all nuclear TDP-43 droplets were of similar structure and composition. For this purpose, we determined the colocalization of endogenous TDP-43 in HEK293 cells with a broad panel of nuclear membraneless organelles (**Figure 5A**). This approach revealed that a fraction of TDP-43 droplets, in particular the largest in size, were labeled with markers of Cajal bodies (**Figure 5A-B**), the maturation compartments of spliceosomal small nuclear ribonucleoprotein particles. Interestingly, TDP-43 was present in all Cajal bodies in HEK293 cells and neurons (**Figure 5C-D**), but Cajal bodies only comprised a small fraction of all TDP-43 droplets. Another portion of TDP-43 was localized within paraspeckles, but, unlike Cajal bodies, not every paraspeckle contained TDP-43. TDP-43 was absent from other RNA-nucleated compartments such as nuclear speckles (*54*) and from protein-exclusive compartments like promyelocytic leukemia (PML) bodies (**Figure 5A-B**) (*55*). In contrast to WT TDP-43, RNA binding-deficient GFP-TDP-43 was incorporated into neither Cajal bodies nor paraspeckles, in line with previous reports (*27*) (**Figure 5E-F and S5A-B**). Intriguingly, also monomeric GFP-TDP-43 was largely absent from these nuclear compartments (**Figure 5E-F and S5A-B**), highlighting the requirement of TDP-43 oligomerization for its incorporation within functional nuclear bodies. Collectively, these observations suggest that at least a fraction of TDP-43 LLPS arises from an RNA scaffold, since TDP-43 has been shown to bind small Cajal body-specific RNAs (*56*) and *NEAT1*, the architectural RNA of paraspeckles (*20, 21, 53*). The remaining, unidentified nuclear TDP-43 droplets, present in both the WT and RNA binding-deficient variant (**Figure 5E-F and S5A-B**), might represent a precursor pool for Cajal bodies and paraspeckles, an inert droplet population or even be linked to yet undefined bodies and functions. Whether any of these potentially “RNAless” TDP-43 nuclear compartments formed by the RNA binding-deficient variant are identical to the unidentified droplets occuring in physiological conditions and/or have functional roles remains unclear. Overall, the heterogeneous nature of nuclear TDP-43 droplets suggests that LLPS is required for a wide array of TDP-43 functions within the nucleus.

**Figure 5.**
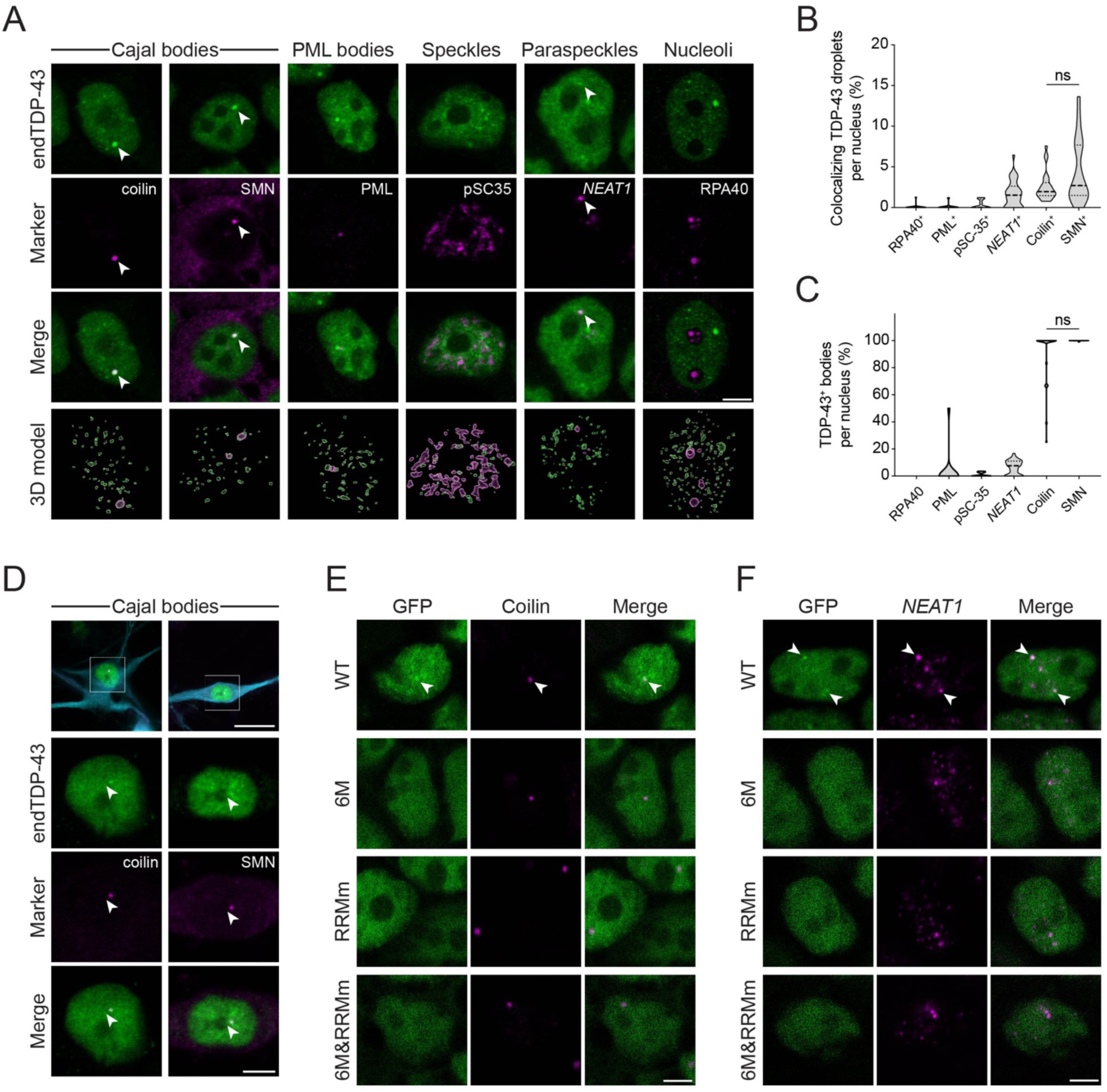
Cajal bodies and paraspeckles are the main TDP-43-containing nuclear bodies. **(A)** Representative confocal microscopy images of HEK293 cells probed for endogenous TDP-43 (endTDP-43) and different subnuclear compartment markers. Lower panel shows the three-dimensional (3D) reconstruction of the endogenous TDP-43 (endTDP-43) droplets and the indicated nuclear body obtained from the confocal Z-stacks. Scale bar: 5 µm. **(B)** Quantification of the 3D reconstructions shown in A depicting the percentage of nuclear TDP-43 droplets colocalizing with markers of subnuclear compartments. N=14-38 cells. **(C)** Quantification of 3D reconstructions shown in A depicting the percentage of each of the analyzed subnuclear compartments that colocalize with endogenous TDP-43. N=14-38 cells. **(D)** Representative confocal microscopy images of human neurons showing the presence of TDP-43 in Cajal bodies. Scale bar: 20 µm (inset: 5 µm). **(E)** Representative confocal microscopy images of the isogenic HEK293 lines expressing the different GFP-TDP-43 variants for 24 h and stained for the Cajal body marker coilin. Scale bar: 5 µm. **(F)** Representative confocal microscopy images of the isogenic HEK293 lines expressing the different GFP-TDP-43 variants for 24 h and hybridized with a fluorescent *NEAT1* probe to mark the paraspeckles. Scale bar: 5 µm.

### TDP-43 oligomerization is required for the transcriptome-wide splicing regulation of its RNA targets

NTD-mediated oligomerization is required for splicing of at least a subset of the RNA targets of TDP-43 (*15, 35*–*38*), but given the broad role of TDP-43 in regulating splicing events (*20, 21, 57, 58*), the question remained whether oligomerization is essential for all its splicing targets. RNA-sequencing (RNA-seq) of our isogenic cell lines revealed that expression of the WT variant resulted in alternative splicing of >70 genes when compared to the expression of the GFP-TDP-43 RRMm, an established splicing-deficient protein version (*16, 18*), including previously reported events of exon inclusion/exclusion, intron retention and alternative polyadenylation site usage depending on TDP-43 binding (*20, 21, 58*) (**Figures 6A and S6A**). When the same comparison was performed between the oligomerization-deficient and the RRMm GFP-TDP-43, no significant alternatively spliced events were found (**Figures 6A and S6A**), suggesting a lack of splicing functionality of monomeric GFP-TDP-43, which was also observable at the differentially expressed RNA and protein levels (**Figure S6B**). A particularly interesting event modulated by TDP-43 binding is the splicing of an alternative intron (intron 7) in its own 3’ UTR, which results in autoregulation of the TDP-43 mRNA and protein levels (*20, 40, 59*) (**Figure S6C**). Analysis of the 3’ UTR of TDP-43 by RNA-seq and qPCR showed that, similar to other alternative splicing events, 6M GFP-TDP-43 cannot promote the exclusion of intron 7 in the endogenous *TARDBP* mRNA (**Figure 6B and S6D-E**), resulting in lack of autoregulation at the protein level (**Figure 6C-D**). This lack of splicing activity by monomeric TDP-43 was not due to its reduced protein concentration, since WT GFP-TDP-43 autoregulated endogenous TDP-43 at comparable expression levels (**Figure S6D-E**). Overall, our data support the requirement of TDP-43 oligomerization for its broad role in splicing regulation.

**Figure 6.**
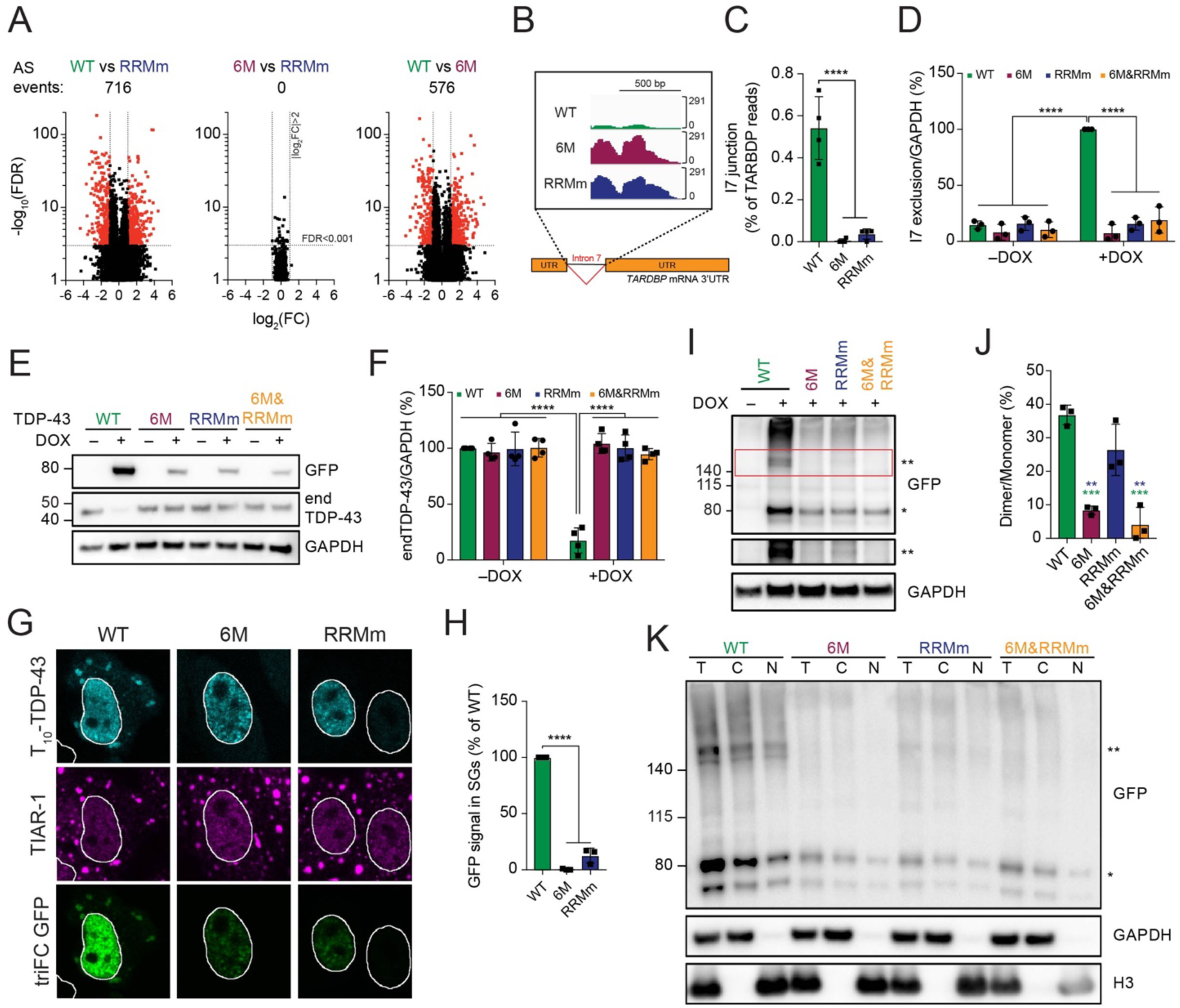
TDP-43 oligomerization is required for splicing regulation in the nucleus and stress granule incorporation in the cytoplasm. **(A)** Volcano plots showing alternative splicing events upon expression of GFP-TDP-43 variants for 48 h. **(B)** RNA sequencing (RNA-seq) coverage across the intron 7 of the *TARDBP* gene, showing a strong decrease in the WT, but not the mutant, GFP-TDP-43-expressing cells. AS events: alternative splicing events. **(C)** Quantification of the RNA-seq reads spanning the intron 7 junction. One-way ANOVA with Tukey’s multiple comparisons post hoc test. **(D)** Endogenous TDP-43 (endTDP-43) intron 7 exclusion levels after expression of the GFP-TDP-43 variants for 48 h in the isogenic cell lines measured by qPCR with primers specifically targeted to the transcripts excluding this region. Two-way ANOVA with Tukey’s multiple comparisons post hoc test. **(E)** Western blot analysis of the isogenic HEK293 after GFP-TDP-43 expression for 48 h showing that only the WT variant regulates endTDP-43 levels. **(F)** Quantification of the endTDP-43 signal from E. N=4 independent experiments. Two-way ANOVA with Tukey’s multiple comparisons post hoc test. **(G)** Tripartite GFP complementation assay using a pair of N-terminally T_10_- and T_11_-tagged TDP-43 constructs co-transfected with GFP_1-9_ in HeLa cells and subjected to arsenite stress for 30 min. TriFC: trimolecular fluorescence complementation. Scale bar: 10 µm. **(H)** Quantification of the signal trimolecular fluorescence complementation (triFC) of GFP in the TIA-1-marked SGs from the immunocytochemistry images shown in G. N=3 independent experiments. Repeated measures one-way ANOVA with Greenhouse-Geisser correction and Tukey’s multiple comparisons post hoc test. **(I)** Expression of GDP-TDP-43 mutNLS variants was induced with doxycycline (DOX) for 4 h before crosslinking protein-protein interactions with DSG and subsequent analysis by western blot. * and ** indicate GFP-TDP-43 monomers and dimers, respectively. **(J)** Quantification of the GFP signal from I. N=3 independent experiments. Repeated measures one-way ANOVA with Greenhouse-Geisser correction and Tukey’s multiple comparisons post hoc test. **(K)** After expression of GFP-TDP-43 mutNLS variants for 48 h, HEK293 cells were treated with DSG to cross-link protein-protein interactions before performing nucleocytoplasmic fractionation and western blot analysis. * and ** indicate GFP-TDP-43 monomers and dimers, respectively. ** p<0.01, *** p<0.001, **** p<0.001. Graph bars represent mean ± SD.

### Cytoplasmic TDP-43 oligomerization is required for its incorporation into stress granules

To understand whether oligomerization is also required for TDP-43 functions outside of the nucleus, we studied TDP-43 incorporation into SGs in the cytoplasm (*29*). Since relocation of TDP-43 into SGs depends on the cell type and the stressor applied (*29*), we resorted to the GFP triFC assay (**Figure S2L**) to investigate TDP-43 complementation in the well-established oxidative stress model in HeLa cells (*29, 30*) T_10_- and T_11_-tagged TDP-43 variants and incubation with recombinant GFP_1-9_ revealed that, unlike the WT protein –but similarly to the RNA binding-deficient mutant–, monomeric TDP-43 did not incorporate into SGs (**Figure 6G-H**), suggesting that TDP-43 oligomerization also takes place in the cytoplasm. To confirm the presence of cytoplasmic TDP-43 oligomers in physiological conditions, we developed isogenic HEK293 Flp-In T-REx lines harboring one copy of each of the GFP-TDP-43 variants in combination with previously published mutations in the nuclear localization signal that abolish nuclear import (GFP-TDP-43 mutNLS) (*60*). Protein levels of the GFP-TDP-43 mutNLS variants were similar to that of their nuclear counterparts (**Figure S6F and 1D**). Interestingly, the localization of the GFP-TDP-43 mutNLS variants differed between the four cell lines. WT and RRMm GFP-TDP-43 mutNLS were predominantly present in the cytoplasm, whereas their monomeric counterparts significantly shifted their localization to the nucleus (**Figure S6G-H**). This suggests that WT and RRMm GFP-TDP-43 mutNLS oligomerize in the cytoplasm, which hinders their passive diffusion back into the nucleus. To confirm this, we performed DSG cross-linking of protein-protein interactions in the GFP-TDP-43 mutNLS lines and found that both WT and RRMm GFP-TDP-43 mutNLS formed oligomers, albeit their presence was reduced in the RRMm variant (**Figure 6I-J**). Moreover, DSG cross-linking followed by nucleocytoplasmic fractionation enabled the detection of WT and RRM GFP-TDP-43 oligomers in the cytoplasmic fraction (**Figure 6K**). Albeit less abundant than nuclear oligomerization, cytoplasmic oligomerization was also observed at endogenous TDP-43 levels by PLA (**Figure 4E**). Altogether, our observations suggest that oligomerization is essential in both the nucleus and the cytoplasm for TDP-43 to perform its functions in RNA metabolism.

### Loss of RNA binding or oligomerization differentially modulate the subcellular localization of TDP-43 inclusions

Decline of the cellular proteostasis capacity with age contributes to protein misfolding in neurodegenerative diseases (*43*), often resulting in the accumulation of ubiquitinated inclusions in affected tissues, including TDP-43 proteinopathies (*2, 3*). Since monomeric and RNA binding-deficient TDP-43 showed shorter half-lives at physiological levels (**Figure 1D-E**), we sought to determine how failure of the UPS machinery affects the accumulation of these species. By blocking the proteasome with the inhibitor MG132, we observed that both monomeric and RNA binding-deficient GFP-TDP-43 formed protein inclusions in the isogenic cell lines, in contrast to the WT counterpart, which remained largely diffuse (**Figure 7A-B and S7A**). The vast majority of aggregates formed by monomeric GFP-TDP-43 (6M and 6M&RRMm) localized to the cytoplasm, in line with previous results showing that high overexpression of oligomerization-deficient GFP-TDP-43 by transient transfection triggers cytoplasmic TDP-43 aggregation (*15*). Interestingly, in addition to cytoplasmic inclusions, MG132 treatment resulted in nuclear aggregation of RNA binding-deficient GFP-TDP-43 in >70% of the cells. The combined loss of oligomerization and RNA binding shifted the aggregation to the cytoplasm, suggesting that nuclear TDP-43 aggregation depended on NTD interactions. The observed TDP-43 aggregation patterns were specific to the inhibition of the UPS degradation pathway, as several classes of proteasome inhibitors, but not an autophagy one, yielded similar outcomes in the isogenic cell lines (**Figure S7B-C**).

**Figure 7.**
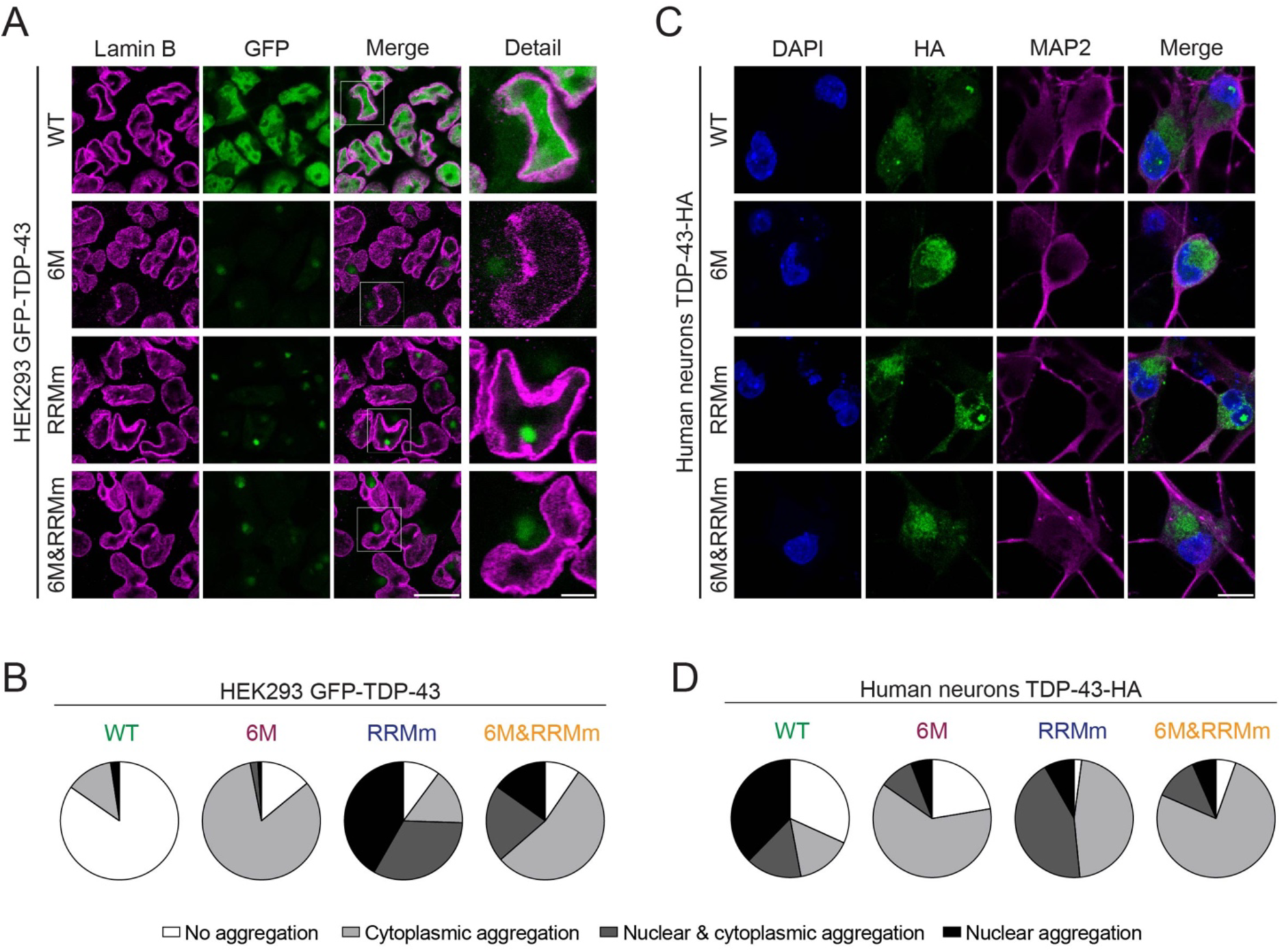
Loss of RNA binding or oligomerization differentially modulate the subcellular localization of TDP-43 inclusions. **(A)** Representative confocal microscopy images of the isogenic GFP-TDP-43 lines after 48 h of expression, treated with the proteasome inhibitor MG132 for the last 24 h and stained for lamin B to mark the nuclear envelope. Note the different localization of TDP-43 inclusions in the oligomerization- (6M and 6M&RRMm, cytoplasmic) versus RNA binding-deficient (RRMm, nuclear) variants. Scale bar: 20 µm (5 µm for inset). **(B)** Quantification of the differentially localized GFP-TDP-43 inclusions after MG132 treatment for the different variants in the isogenic HEK lines as shown in A. Represented values are averages from N=3 replicates, with N=189-497 cells quantified per condition and replicate. **(C)** Representative maximum intensity Z-projections from confocal fluorescence imaging (thickness of 4 µm, in steps of 1 µm) of human neurons transduced with TDP-43-HA variants and treated overnight with the proteasome inhibitor MG132. Scale bar: 10 µm. **(D)** Quantification of the differentially localized TDP-43-HA inclusions in human neurons as described in C. Represented values correspond to the quantification of N=85-97 neurons from two independent experiments.

In human neurons, monomeric TDP-43-HA variants also predominantly aggregated in the cytoplasm, whereas RRMm TDP-43-HA additionally presented nuclear inclusions in >50% of transduced neurons (**Figure 7C-D**), thus reproducing the distinct TDP-43 aggregation patterns observed in isogenic cell lines. Upon MG132 treatment, WT TDP-43-HA also formed inclusions in neurons, both in the nucleus and cytoplasm, likely due to higher transgene protein levels in transduced neurons compared to the isogenic GFP-TDP-43 lines. Interestingly, and in line with higher protein levels, RRMm TDP-43-HA already formed nuclear inclusions in the absence of proteasome inhibition in a subset of transduced human neurons (**Figure S7D**). Taken together, our data show that loss of oligomerization shifts TDP-43 aggregate formation from the nucleus to the cytoplasm.

### TDP-43 aggregates in an LLPS- or an aggresome-dependent manner in the nucleus and cytoplasm, respectively

To understand the origin of cytoplasmic and nuclear TDP-43 inclusions, we performed live cell imaging of the GFP-TDP-43 isogenic lines during the treatment with the proteasome inhibitor. While WT GFP-TDP-43 droplets merely changed position, fused and split within the nucleus upon MG132 addition, monomeric GFP-TDP-43 formed a single cytoplasmic inclusion whose size increased over time, accompanied by gradual decrease in the diffused nuclear TDP-43 signal (**Figure 8A**) resembling the nuclear clearance that has been widely reported in neurons with TDP-43 pathology in ALS and FTLD patients (*1*–*3*). Fluorescence recovery after photobleaching (FRAP) experiments revealed that whereas WT GFP-TDP-43 remained diffuse throughout the treatment, cytoplasmic inclusions comprising monomeric TDP-43 were immobile structures, (**Figure 8B and S8A-B**). A similar aggregation pathway –a single focus expanding in size yielding one solid cytoplasmic inclusion– was also observed for the RRMm GFP-TDP-43 in the cytoplasm (**Figure S8C-D**), suggesting that the formation of cytoplasmic aggregates upon proteasomal failure does require neither oligomerization nor RNA binding. In contrast, in the nucleus, the elevated protein levels of RRMm GFP-TDP-43 caused by proteasome inhibition induced the formation of many initially dynamic droplets, which eventually fused to form a single solid inclusion (**Figure 8B-C**), reminiscent of the single intranuclear inclusions found in patients with specific FTLD subtypes (*6, 61*). FRAP analysis showed that GFP-TDP-43 RRMm in the final inclusion had lost its fluid behavior (**Figure 8B**). In a subset of cells, RRMm GFP-TDP-43 deposited in the nucleoli (**Figure S8C**), in line with the protein quality control properties of this phase-separated compartment (*62*). Importantly, also in human neurons the formation of nuclear aggregates in a subset of transduced cells expressing RRMm TDP-43-HA (**Figure S7B**) was accompanied by the pronounced presence of nuclear droplets in transduced cells without inclusion (**Figure S8F**). Together, these data suggest that TDP-43 aggregates via LLPS in the nucleus.

**Figure 8.**
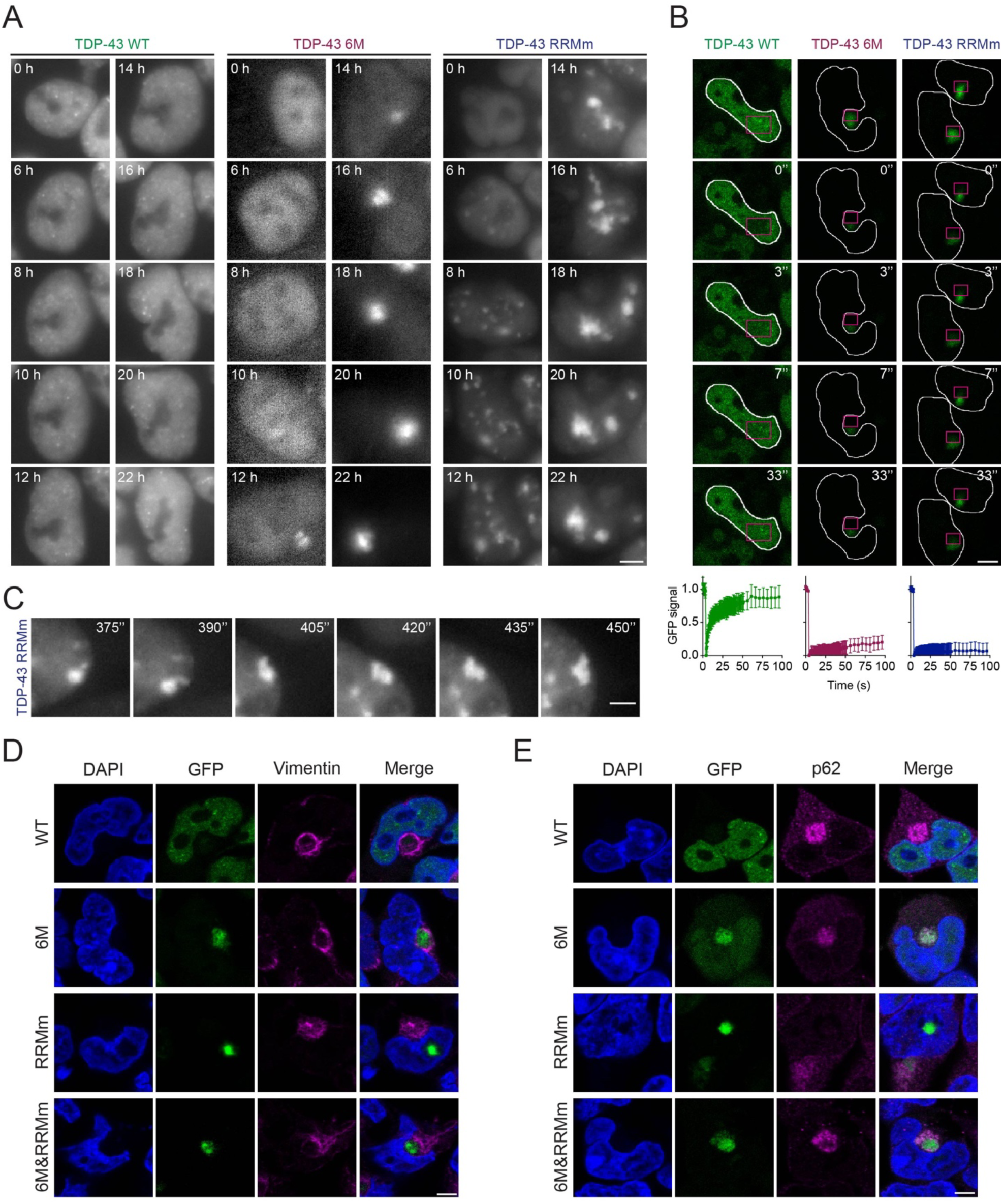
TDP-43 aggregates in an LLPS- or an aggresome-dependent manner in the nucleus and cytoplasm, respectively. **(A)** Representative images of live widefield fluorescence microscopy over the course of the MG132 treatment of the isogenic GFP-TDP-43 lines in the conditions described in Figure 7A. Numbers in images indicate the experimental time point in hours (h) of MG132 treatment. Scale bar: 5 µm. **(B)** Representative fluorescent confocal microscopy images of FRAP experiments in the GFP-TDP-43 aggregates originated upon MG132 treatment as described in Figure 7A. FRAP was performed in the areas highlighted in magenta. Bottom panel: Measured GFP values are expressed as a fraction of the average pre-bleach fluorescence levels. Scale bar: 5 µm. **(C)** Live widefield fluorescence microscopy depicting a fusion event and aberrant phase transition of RRMm GFP-TDP-43 droplets upon MG132 treatment in the conditions described in Figure 7A. Scale bar: 3 µm. **(D)** Representative confocal microscopy images of the isogenic GFP-TDP-43 lines in the conditions described in figure 7A and stained for vimentin. Scale bar: 5 µm. **(E)** Representative confocal microscopy images of the isogenic GFP-TDP-43 lines in the conditions described in Figure 7A and stained for p62. Scale bar: 5 µm.

Cytoplasmic TDP-43 inclusions were consistently found adjacent to the nucleus, in a location occupied by the aggresome, a juxtanuclear accumulation of misfolded proteins resulting from saturation of the chaperone refolding system and/or the UPS degradation pathway (**Figure S8G**) which has been linked to the origin of protein aggregates in neurodegenerative diseases (*63*). Indeed, cytoplasmic inclusions, formed by 6M, RRMm or 6M&RRMm GFP-TDP-43, were surrounded by a vimentin cage, a characteristic feature of aggresomes (**Figure 8D and S8H**) (*63*). Additionally, cytoplasmic, but not nuclear, TDP-43 aggregates were positive for p62 (**Figure 8E**), a critical component of aggresomes (*64*) and a pathological aggregate marker in certain FTLD subtypes (*6, 61*). These observations suggest that distinct pathways towards TDP-43 aggregation are at play in the nucleus and cytoplasm and that monomerization increases TDP-43 incorporation in cytoplasmic aggresomes upon proteasomal failure, thereby potentially triggering cytoplasmic TDP-43 aggregation in disease.

## Discussion

In this study, we describe the interconnection between NTD-driven TDP-43 oligomerization and RNA binding, and show that they cooperatively retain TDP-43 in the nucleus, instruct its functionality and therefore slow down its turnover. We demonstrate that oligomerization is essential for the broad function of TDP-43 in splicing regulation and allows its LLPS-mediated incorporation into nuclear membraneless compartments, including Cajal bodies and paraspeckles. Our work describes for the first time that, under physiological conditions, TDP-43 oligomers exist in the cytoplasm, where they are required for LLPS-dependent incorporation into SGs. Moreover, we show that, in the absence of RNA binding, TDP-43 oligomerization is reduced, but the dimers that do form adopt a spatial conformation that is different from the RNA-bound oligomeric state. Importantly, our results shed light on the molecular mechanism of two distinct and independent pathways triggering TDP-43 aggregation, which highlight the importance of TDP-43 monomerization and/or loss of RNA binding as key early events in the development of TDP-43 proteinopathies (**Figure 9**).

**Figure 9.**
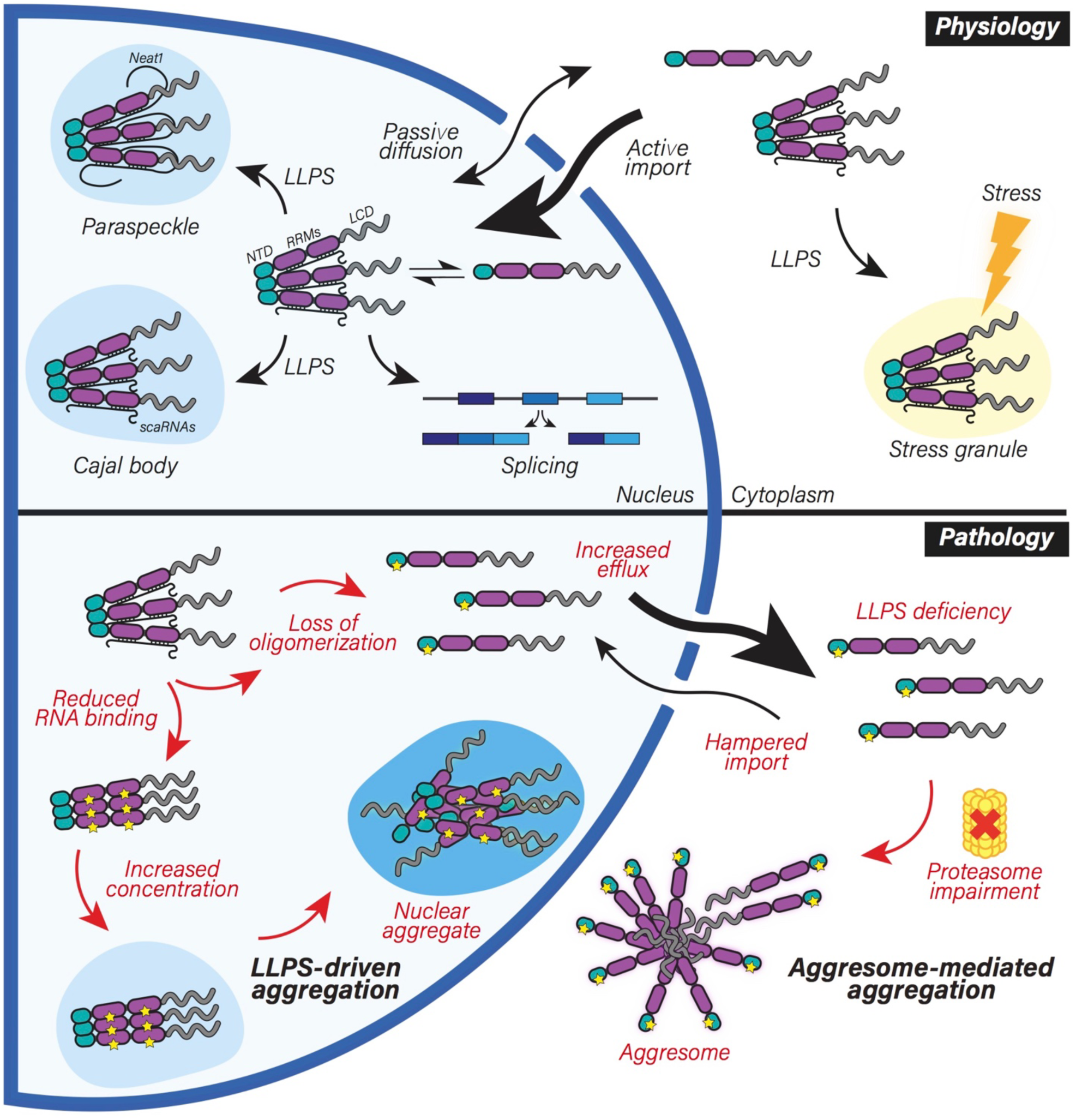
Oligomerization and RNA-binding enable TDP-43 physiological functions and their disruption drives nuclear and cytoplasmic aggregate formation via distinct pathways. Schematic representation of the role of NTD-driven oligomerization and RNA binding in TDP-43 physiology and pathology. **Upper panel:** In healthy cells, TDP-43 monomers and oligomers are in a dynamic equilibrium. TDP-43 is actively imported into the nucleus (*12*), where oligomerization and RNA binding retain it in large macromolecular complexes, limiting passive outflow. In the nucleus, oligomerization and RNA binding are key for the essential roles of TDP-43 in RNA processing, including alternative splicing. Furthermore, oligomerization enables the LLPS of TDP-43 and in conjunction with binding to specific RNAs –such as small Cajal body-specific RNAs (scaRNAs) (*56*) and *NEAT1* (*20, 21*)– allows its localization to distinct subnuclear compartments, primarily Cajal bodies and paraspeckles. TDP-43 oligomers are also detected in the cytoplasm, where its LLPS-mediated incorporation into SGs under stress conditions depends on both oligomerization and RNA binding. **Lower panel:** In disease, loss of TDP-43 oligomerization or RNA binding increases the nuclear efflux of LLPS-deficient monomers, disrupts its inclusion into nuclear bodies, leads to transcriptome-wide splicing alterations (including TDP-43 auto-regulation) and drives TDP-43 aggregation via two independent pathways. Upon failure of the ubiquitin-proteasome degradation machinery observed with aging and in ALS/FTLD patients (*2, 3, 43*), monomeric TDP-43 aggregates in an aggresome-dependent manner in the cytoplasm. Notably, the known decline in active nuclear import in disease (*83*) would further exacerbate TDP-43 cytoplasmic accumulation. In the nucleus, loss of TDP-43 RNA-binding results in enhanced formation of both monomers that escape to the cytoplasm and conformationally distinct TDP-43 oligomers. When the concentration rises (e.g. due to the aforementioned proteasomal failure), RNA-binding deficiency modulates TDP-43 LLPS, culminating in the formation of nuclear aggregates via an LLPS-mediated pathway. Taken together, RNA binding and oligomerization allow TDP-43 to maintain its localization and function in physiology and their disruption drives LLPS-dependent and aggresome-dependent aggregation pathways in the nucleus and cytoplasm, respectively. LCD: low complexity domain, LLPS: liquid-liquid phase separation, NTD: N-terminal domain, RRMs: RNA recognition motifs.

TDP-43 oligomerization is a dynamic event that controls the relative amounts of TDP-43 monomers, dimers and oligomers in the cell. However, the specialized roles of the individual TDP-43 species in health and disease remained unknown. Expression of different TDP-43 variants at near-physiological levels in our human cell line and neural system revealed that, in comparison to its oligomeric counterpart, monomeric TDP-43 lacks functionality, becomes more prone to escape the nucleus –likely by passive diffusion (*32, 47*)– and is rapidly degraded. In the event of proteasomal failure, these otherwise short-lived TDP-43 monomers are deposited into aggresomes, whose expansion is accompanied by a progressive decrease in nuclear TDP-43. This cytoplasmic aggregation observed in our human cellular models recapitulates the key pathological TDP-43 features observed in affected neurons of patients with TDP-43 proteinopathies, namely its nuclear clearance, loss of function and cytoplasmic aggregation (*2, 3*). Interestingly, the aggresome pathway was previously linked to sporadic ALS/FTLD (*65*) through the disturbance of p62 (*65, 66*), a key player in aggresome formation (*64*). Furthermore, aggresome markers HDAC6 and p62 have been reported to colocalize with a subset of cytoplasmic TDP-43 aggregates in ALS/FTLD patients (*6, 61, 67*). When the conditions that favor the monomeric state of TDP-43 and hamper its proteasomal degradation persist, TDP-43 monomers contained in the aggresomes may resist clearance by aggrephagy and rather further mature into compact aggregates due to the high concentration of unfolded LCRs in the absence of NTD-driven organization that keeps them apart (*15*). Since monomeric TDP-43 is unable to autoregulate its own levels, the continued production of more TDP-43 to compensate for its loss of function will only exacerbate this pathological transition. Our data signifies that loss of TDP-43 oligomerization ignites a pathological cascade that culminates in the formation of cytoplasmic TDP-43 inclusions via the aggresome pathway.

In addition to loss of oligomerization, our data indicate that also the disturbance of TDP-43 oligomer conformation triggered by loss of RNA binding plays a role in pathology through a distinct molecular pathway in the nucleus. While previous studies have addressed the role of RNA in TDP-43 pathogenesis, our work provides insights into the molecular mechanism underlying the aberrant phase transition of NTD-driven RNAless TDP-43 oligomers into nuclear immobile inclusions. Using a combination of imaging and biochemical assays, we observed that TDP-43 oligomers present a different conformation in an RNA-bound or - unbound state. RNA-bound TDP-43 oligomers enact its physiological functions, maintain its localization and antagonize the formation of pathological aggregates (*15* and this study), while RNAless TDP-43 oligomers, that may or may not have functional roles, undergo aberrant phase separation leading to nuclear aggregation (*37* and this study). These observations clarify the apparently controversial role of the NTD in aggregate formation, which has been found both synergistic (*35, 68*–*70*) and antagonistic (*15*). Based on our observations, we propose that NTD-mediated TDP-43 self-interaction is a double-edged sword: in the presence of RNA, it is essential for TDP-43 to perform its functions and undergo physiological LLPS, while in the absence of RNA binding it promotes aberrant LLPS that leads to aggregation.

In addition to the cytoplasmic aggregation of TDP-43, neuronal intranuclear inclusions have been reported in two of the five recognized FTLD-TDP subtypes (*2, 3, 6, 71*–*73*) and are particularly abundant in FTLD cases linked to mutations in valosin-containing protein (VCP) (*71, 72*), which is involved in nuclear protein quality control degradation (*74*). In our human cellular models, we observed LLPS-driven intranuclear aggregation of RNA binding-deficient TDP-43 oligomers upon inhibition of proteasomal degradation. Similarly, nuclear inclusions have also been reported in previous studies employing strong overexpression of TDP-43 RNA binding mutants in cells and neurons (*23, 33, 36, 42, 60*). In addition to its LLPS-mediated nuclear aggregation, RNA binding-deficient TDP-43 formed cytoplasmic inclusions via the aggresome pathway. This route is likely favored by the increase of the monomeric state in the absence of RNA binding, accompanied by nuclear efflux, as previously shown upon proteasome inhibition in cultured neurons (*75*). Altogether, our data show that when a cell encounters proteostatic stress, TDP-43 takes different routes towards inclusion formation, and the selection of the pathway depends on both the protein state (monomeric versus RNAless) and the subcellular environment (cytoplasm versus nucleus). While our study provides evidence for the importance of two such routes (nuclear LLPS-dependent and cytoplasmic aggresome-mediated), additional aggregation pathways –for example, cytoplasmic LLPS-mediated (*24, 33, 67*)– likely exist and may be triggered under different circumstances and involve other TDP-43 states. Collectively these distinct pathways may account for the spectrum of cytoplasmic aggregates observed in patients with TDP-43 proteinopathies.

Why would the majority of TDP-43 aggregates reside in the cytoplasm in ALS/FTLD patients? Our data indicates that TDP-43 monomerization and its subsequent nuclear efflux is a more frequent or potent event than decreased RNA binding affinity in these cases. Alternatively, the high nuclear RNA concentrations that have been shown to prevent LLPS of RNA-binding proteins (*50*) may counteract aberrant LLPS and aggregation, even in the absence of specific RRM-RNA interactions for TDP-43. This balancing mechanism may be reinforced by the upregulation of some architectural TDP-43 RNA targets in the nucleus, including *NEAT1* (*26, 50*). This binding could instruct the proper, RNA-loaded orientation of oligomers with a concomitant increase in physiological LLPS in the form of nuclear bodies. In fact, elevated *NEAT1* levels and paraspeckle formation have been amply reported in TDP-43 proteinopathies (*21, 26, 76, 77*).

The cellular machinery that regulates TDP-43 oligomerization remains unknown. While protein concentration is a determinant of TDP-43 oligomerization (*14, 15*), the conditions that keep the balance between monomeric and oligomeric TDP-43 species in healthy cells and, most importantly, increase TDP-43 monomerization in disease require further investigation. Post-translational modifications (PTMs) are excellent candidates for such physiological regulation of TDP-43 oligomerization and a recent study identified a single serine phosphorylation event within the NTD interface that decreases oligomerization *in vitro* (*38*), albeit its effects within cells or occurrence in disease have not yet been tested. Similarly, specific acetylation events within the RRMs that lower the affinity of TDP-43 for RNA have been detected in ALS patients (*78*) and were subsequently reported to trigger nuclear TDP-43 aggregation via aberrant LLPS (*34*). Supporting the link between reduced RNA affinity and disease, ALS/FTD-associated mutations within the RRMs have been shown to disrupt RNA binding and enhance TDP-43 proteinopathy (*79*). Moreover, since ATP was recently shown to directly bind the NTD of TDP-43, thereby enhancing its oligomerization (*44*), the decrease in cellular ATP levels with age (*80*) could also act as a monomerization-inducing trigger. It is also conceivable that additional, yet unknown, protein interactors of TDP-43 may act as modifiers of oligomerization. Future work should focus on determining the molecular switches within TDP-43 and/or protein partners that regulate its oligomerization and RNA binding and on validating the occurrence of diminished TDP-43 oligomerization and RNA affinity in patient tissue. Such insights will be valuable to inform drug design targeting TDP-43 oligomerization and RNA binding, including dimer stabilization or recovery of RRM-RNA interactions, among others.

In conclusion, oligomerization and RNA binding allow TDP-43 to maintain its localization and function in physiology and their disruption drives distinct aggregation pathways in the nucleus and cytoplasm, indicating that distinct molecular origins may account for the plethora of TDP-43 aggregation types observed in ALS and FTLD subtypes.

## Materials and Methods

### Plasmids

The pcDNA5 plasmid containing the GFP-tagged human TDP-43 cDNA sequence was a kind gift of Dr. Shuo-Chien Ling (*41*). The mutations to introduce a siRNA resistance in the TDP-43 coding region without altering the amino acid sequence and the oligomerization-disrupting mutations in the N-terminal domain were previously described (*15*). RNA-binding disruption mutations were introduced by site-directed mutagenesis PCR as described before (*15*), using high fidelity Phusion DNA Polymerase (New England Biolabs, M0530) with primers detailed in **Table S1** followed by DpnI (New England Biolabs, R0176) digestion before bacterial transformation and colony selection.

Plasmids encoding T_10_- and T_11_-tagged TDP-43 and TDP-43 6M have already been reported (*15, 52*). The sequence encoding TDP-43 RRMm was amplified by PCR from the pcDNA5 plasmids described above by adding BamHI/XhoI restriction sites using the primers detailed in **Table S1**. Amplified products were cloned into the T_10_ and T_11_pcDNA3 parental plasmids

(*52*) between BamHI and XhoI in order to obtain plasmids encoding T_10_-HA-TDP43 RRMm and T_11_-β1-TDP-43 RRMm. The same amplified sequence was cloned into the plasmids encoding T_10_- and T_11_-tagged TDP-43 6M (*52*) between EcoRI and XhoI in order to obtain plasmids encoding for T_10_-HA-TDP43 6M&RRMm and T_11_-β1-TDP-43 6M&RRMm.

TDP-43 with a C-terminal HA tag was directly amplified from the GFP-TDP-43 construct described above and inserted into an autoregulatory all-in-one TetON cassette previously inserted into pLVX backbone (*81*) by Gibson cloning using the NEBuilder HiFi DNA Assembly Cloning Kit (New England Biolabs, E5520S) according to the manufacturer’s instructions.

The plasmid encoding the His6-tagged RRMs of TDP-43 for bacterial protein expression was previously published (*16*) and mutations disrupting RNA binding were introduced as described above. The plasmid encoding TDP-43-MBP-His6 for bacterial protein expression was a gift from Nicolas Fawzi (Addgene plasmid #104480; http://n2t.net/addgene:104480; RRID:Addgene_104480) (*38*). Point mutations for 6M, S2C, C39S and C50S were introduced with site-directed mutagenesis PCR with high fidelity Phusion DNA Polymerase (New England Biolabs, M0530) using primers described in **Table S1**. The plasmid encoding the MBP-His6 was generated using deletion cloning by PCR with high fidelity Phusion DNA Polymerase (New England Biolabs, M0530) using designed primers described in **Table S1**.

### Recombinant protein expression and purification

Production of TDP-43 RRMs for NMR studies was performed as previously described (*16*). Production of full-length TDP-43 was performed as previously reported (*7*). For more details, see **Supplementary materials & methods**. The MBP-His6 was purified as the full-length TDP-43 but as purity was reached after the Ni Sepharose Excel material the protein was subsequently dialysed against a storage buffer (20 mM Tris pH 8.0, 300 mM NaCl, 1 mM TCEP) overnight at 4°C, concentrated the next day using Amicon Ultra-15 concentrators MWCO 10 kDa (Merck Millipore, UFC901024), flash frozen and kept at -80°C.

### Circular dichroism experiments

TDP-43-MBP constructs were thawed on ice and centrifuged for 15 min at 17 100 *g* and 4°C. The buffer was exchanged to 10 mM sodium phosphate pH 7.4 (0.036 % (w/v) sodium phosphate monobasic (Sigma Aldrich, S3139), 0.2 % (w/v) sodium phosphate dibasic (Sigma Aldrich, S5136)) using 500 µl concentrators with a MWCO of 10 kDa (Merck Millipore, UFC501024) 5 x (11 000 *g*, 10 min, 4°C). To ensure the solubility of the isolated protein, the concentrate was again centrifuged as described above. Protein concentration was determined by the extinction coefficients and molecular weights of the constructs (ExPASy ProtParam software) and absorbance at 280 nm (NanoDrop, Thermo Scientific). CD spectra of 200 µg/ml recombinant protein were recorded at 20 °C from 180-250 nm with a bandwidth of 1 nm and a sampling period of 25 s on a Chirascan V100 (Applied Photophysics). Ellipticity (in mdeg) was transformed to molar ellipticity (in deg cm^2^ dmol^− 1^) with the following formula, where c (in M) is the concentration and L (in cm) represents the pathlength of the cuvette:

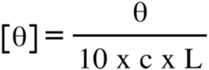

### Phase separation experiments

Pure concentrated fractions of recombinant protein MBP-tagged full-length WT and 6M TDP-43 were desalted in a buffer of 20 mM sodium phosphate pH 7.2 (0.072 % (w/v) sodium phosphate monobasic, 0.4 %(w/v) sodium phosphate dibasic), 300 mM NaCl, 0.001% (v/v) TWEEN 20, 50 mM L-arginine (Merck Millipore, A5006), 50 mM L-glutamic acid (Merck Millipore, G1251) on a HiTrap desalting column (Cytavia, 17140801). An equimolar fluorophore labeling reaction was set up with CF660R maleimide (Biotum Inc., Fremont, CA) previously dissolved in dimethylformamide (DMF) in an N2-hood and incubated for 16 h at 4°C under constant rotation. Reaction was stopped with 10 mM DTT and subsequently passed through a HiLoad 10/300 Superdex 200 pg SEC (Cytiva, 28990944) in SEC buffer on an Äkta pure system (Cytavia). Labelling position was confirmed by mass spectrometry and labelling ratio was determined by first determining the correct protein concentration:

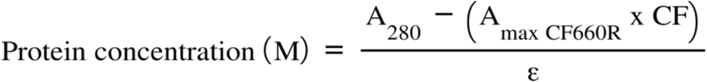

where A_280_ is the absorbance of the protein at 280 nm, A_max CF660R_ is the CF660R dye absorbance at 663 nm (absorbance maximum of CF660R), CF is the correction factor for the amount of absorbance at A_280_ caused by the dye and ε is the protein molar extinction coefficient.

The labelling ratio was subsequently calculated with following formula:

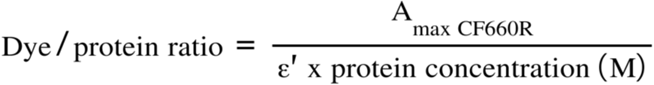

where, ε’ is the molar extinction coefficient of the CF660R fluorescent dye.

Correctly labelled protein with a labeling efficiency of >75% was snap frozen and stored at -20°C. TDP-43-MBP constructs were thawn and centrifuged (17100 *g*, 15 min, 4°C) before exchanging the buffer to a to phase separation buffer consisting of 20 mM HEPES pH 7.4 (Biosolve, 0008042359BS), 150 mM NaCl, 1 mM TCEP. The samples were centrifuged 5 x (11 000 *g*, 10 min, 4°C) and protein concentration determined as described above. Phase separation of 10 µM TDP-43-MBP containing CF660R-labelled TDP-43-MBP at a ratio of 1/200 was induced using dextran (Sigma, D8906) at a final concentration of 10% (w/v) in phase separation buffer and incubation for 2 h at 22°C in µ-Slide Angiogenesis glass-bottom coverslips (ibidi, 81507). Images were acquired on a fluorescence microscope (Nanoimager S, ONI) with an Olympus 100x objective (1.4 NA) using a wavelength of 640 nm and 13% laser power. Reversibility was tested by adding 1,6-hexanediol (Sigma, 240117) to a final concentration of 8% (w/v) in phase separation buffer for 10 min at 22°C and subsequent imaging. Image analysis and droplet counts were performed with ImageJ (1.52k).

### Cell culture

All cells were cultured at 37ºC with saturated humidity and an atmosphere of 5% CO2. HeLa cells were cultured in DMEM (Gibco, 61965-059) supplemented with 1% non-essential amino acids (Gibco, 11140035) and 10% fetal bovine serum (FBS; Gibco, 10270-106).

Mouse motor neuron-like hybrid cells NSC-34 (Bioconcept, CLU140) were proliferated on Matrigel (Corning, 354234)-coated dishes in Dulbecco’s modified Eagle medium (DMEM; Sigma, D5671) supplemented with 10% FBS, 1X GlutaMAX (Gibco, 35050-061), 100 U/ml penicillin and 100 µg/ml streptomycin (Gibco, 15140-122). For experiments, differentiation was induced by switching DMEM/F12 medium (Gibco, 21331-020) supplemented with 1X GlutaMAX, 1X B27+ supplement (Gibco, 17504-044), 1X N2 supplement (Gibco, 17502-048), 100 U/ml penicillin and 100 µg/ml streptomycin.

HEK293 Flp-In T-REx stable lines were cultured in DMEM (Sigma, D5671) supplemented with 10% FBS, 1X GlutaMAX, 15 µg/ml blasticidin S (Gibco, R21001; Invivogen, ant-bl-10p), 100 µg/ml hygromycin B (Gibco, 10687010), 100 U/ml penicillin and 100 µg/ml streptomycin. For details regarding live development, see **Supplementary materials and methods**. When required, cells were treated with the appropriate chemicals at the following concentrations: 1 µg/ml doxycycline (Clontech, 631311), 2.5 µM MG132 (APExBIO, A2585), 1.25 µM MLN9708 (Selleck Chemicals, S2181), 500 nM bortezomib (BTZ; APExBIO, A2614), 100 nM bafilomycin A1 (Sigma-Aldrich, SML1661), 100 µg/ml cycloheximide (Sigma-Aldrich, C4859), 5 µg/ml actinomycin D (Sigma-Aldrich, A1410), 10% 1,6-hexanediol, 1mM sodium arsenite (Sigma-Aldrich, 35000-1L-R), dimethyl sulfoxide (DMSO; Sigma-Aldrich, D2650).

Human neural networks were differentiated from an in-house-developed iPSC-derived self-renewing human neural stem cell line (iCoMoNSCs) obtained from control human skin fibroblast, as described previously (*81*). For details regarding the culture and differentiation, see **Supplementary materials and methods**. When required, neural cultures were treated with the appropriate chemicals at the following concentrations: 1 µg/mL doxycycline, 10 µM MG132, 10 µg/ml actinomycin D, 25 µM ivermectin (Sigma-Aldrich, I8898).

### Lentiviral vector production

TDP-43-HA variants were packaged into lentivirus, harvested and concentrated as described previously (*81*). Lentiviral pellets were resuspended in neural maturation medium (containing all supplements but forskolin and cAMP, see **Supplementary materials and methods**), achieving 10x concentrated preparations of which the lentiviral titer was determined using Lenti-X GoStix Plus (Takara, 631280). Lentiviral preparations were aliquoted and stored at -80°C until use.

### Quantitative PCR (qPCR)

HEK stable lines were plated at a density of 3 × 105 cells/well in a 6-well plate (TPP, 92406). Expression of GFP-TDP-43 was induced with 1 µg/ml doxycycline after 24 hours and cells were harvested 48 h later. Total RNA from cells in a single well of a 6-well plate was isolated using the RNeasy Plus Mini Kit (Qiagen, 74134) according to the manufacturer’s instructions. Complementary DNA (cDNA) corresponding to 1 µg was generated using oligo(dT)20 primers and the SuperScript™ III First-Strand Synthesis SuperMix (Invitrogen, 18080400) according to the manufacturer’s instructions. A qPCR of 50 cycles was performed with 10 ng of cDNA and 6.6 pmol of each of the primers (**Table S2**) per reaction using Fast SYBR™ Green Master Mix (Applied Biosystems, 4385612) in a AriaMx Real-Time PCR System (Agilent, G8830A). Relative fold gene expression was calculated with the 2–ΔΔCt method.

### GFP complementation assay

Tripartite GFP complementation experiments were performed as described before (*15, 52*). For more details, see **Supplementary materials and methods**.

### Protein-protein cross-linking in cells

Native protein-protein interactions were stabilized by crosslinking with disuccinimidyl glutarate (DSG; Thermo Scientific, 20593 or A35392) as previously reported (*15*). In brief, cells grown to 80% confluency in 6-well or 10 cm plates were washed once in cell culture-grade PBS (Gibco, 10010-015), scrapped in ice-cold cell culture-grade PBS and collected at 300 *g* and 4°Cfor 5 min in a 1.5 ml microfuge tube. Cells were resuspended in 100/600 µl ice-cold cell culture-grade PBS containing protease inhibitors and a freshly prepared 20 mM or 100 mM DSG solution in DMSO was added to the suspension to a final concentration of 1 mM. After incubation at 25 ºC and 1500 rpm for 30 min in a Thermomixer (Eppendorf, 2230000048), the reaction was quenched by addition of Tris base to a final concentration of 20 mM and further incubation for 15 min. Cells were collected by centrifugation at 300 *g* for 5 min.

### Nucleocytoplasmic fractionation of cells

Nucleocytoplasmic fractionation of the GFP-TDP-43 (mutNLS) isogenic HEK293 lines was performed following a previously published protocol (*82*) with the following changes. Cells in 10-cm dishes at 80-90% confluency were scrapped in 1 ml cell culture-grade PBS, collected by centrifugation at 300 g for 5 min and resuspended in 880 µl. The final pellet fraction containing the isolated nuclei was resuspended in 380 µl of 1X Laemmli buffer and 1X reducing agent (Invitrogen, B0009) in cell culture-grade PBS supplemented with protease inhibitors, and 10 µl of each three of the final fractions were loaded onto the polyacrylamide gel for western blot analysis.

### Immunocytochemistry

For immunocytochemistry experiments, cell lines were plated onto poly-D-lysine (Sigma-Aldrich, P6407)-coated 96- or 24-well plates (Greiner Bio-One, 655090; ibidi, 82426) or 8-well glass chambers (ibidi, 80827). Unless stated otherwise, cell line cultures were fixed in 4% methanol-free formaldehyde (Thermo Scientific, 28908) in warm medium for 15 min, washed with cell culture-grade PBS, permeabilized and blocked in 10% donkey serum (Sigma-Aldrich, S30-M) and 0.1% Triton X-100 in cell culture-grade PBS for 10 min at RT. Neural cultures were plated onto Matrigel-coated 96-well plates (135 000 cells/well) or 8-well glass-bottom chambers (between 240 0000 and 425 000 cells/well). Neural cultures were fixed in 4% methanol-free formaldehyde in cell culture-grade PBS for 25 min, washed with cell culture-grade PBS, permeabilized in 0.5% Triton X-100 in cell culture-grade PBS for 5 min and blocked in 10% donkey serum and 0.1% Triton X-100 in cell culture-grade PBS for 30 min at RT. Primary antibodies (**Table S3**) were diluted in the same buffer and incubated with the samples overnight at 4°C. After three washes with cell culture-grade PBS, Alexa Fluor-conjugated donkey secondary antibodies (**Table S3**) were incubated for 1h at RT and subsequently further washed with cell culture-grade PBS. Nuclei were counterstained with 1 µg/ml DAPI (Thermo Scientific, 62248) and samples imaged in cell culture-grade PBS.

### Proximity ligation assay (PLA)

Cell lines plated on 96-well plates were fixed in pure ice-cold methanol for 7 minutes at - 20ºC, followed by three cell culture-grade PBS washes. PLA assay was performed using custom labelled antibodies and the Duolink In Situ Detection Reagents Red (Sigma-Aldrich, DUO92008) as described by the manufacturer but with slight modifications. In brief, antibodies of choice (anti-GFP and anti-HA in **Table S3**) in a carrier-free buffer were labelled with MINUS and PLUS probes using the Duolink In Situ Probemaker MINUS (Sigma-Aldrich, DUO92010) and Duolink In Situ Probemaker PLUS (Sigma-Aldrich, DUO92009) according to the manufacturer’s instructions. Cells grown in 96-well plates were fixed in pure ice-cold methanol for 7.5 min at -20°C, followed by two washes with cell culture-grade PBS. After blocking for at least 15 min with blocking solution at room temperature, probe-conjugate antibodies were incubated overnight at 4°C. After two washes of 5 minutes each with buffer A (10 M Tris, 150 mM NaCl 0.05% Tween 20), ligation reaction was performed as indicated by the manufacturer. After an additional two washes of 5 min each with buffer A, amplification reaction was performed as indicated by the manufacturer. The reaction was quenched with buffer B (100 mM Tris, 100 mM NaCl). If required, secondary antibodies were incubated in cell culture-grade PBS overnight at 4°C, followed by DAPI counterstaining and imaging in cell culture-grade PBS.

### Fluorescence in situ hybridization (FISH)

FISH assay was performed using target probes for NEAT1 (Invitrogen, VX-01) and the View RNA Cell Plus Assay (Invitrogen, 88-19000) according to the manufacturer’s instructions with slight modifications. HEK stable lines were plated at a density of 10 × 10^4^ cells/well onto poly-D-lysine-coated 96-well plates. After 24 h, expression of GFP-TDP-43 was induced and, after further 48 h, cells were fixed in ViewRNA Cell Plus Fixation/Permeabilization Solution for 30 min at room temperature. After five washes with 1x PBS with RNase Inhibitor, ViewRNA Cell Plus Probe Solution containing the target probes was incubated for 2 h at 40°C. Following, the cells were washed with ViewRNA Cell Plus Wash Buffer Solution and Signal amplification was continued as indicated by the manufacturer. Nuclei were counterstained with DAPI and samples were imaged in cell culture-grade PBS.

## Acknowledgments

We thank Prof. Benjamin Schuler (Department of Biochemistry, University of Zurich) for critical feedback on this work and Dr. Matthias Gstaiger (Institute of Molecular Systems Biology, ETH Zurich, Switzerland) for sharing the Flp-In T-REx HEK293 cell line and the pOG44 plasmid. We gratefully acknowledge the support from the Proteomics team at the Functional Genomics Center Zurich (FGCZ) from the University of Zurich (UZH), especially Dr. Paolo Nanni, Dr. Serge Chesnov and Dr. Witold Wolski. We also like to thank Dr. Jana Doehner, Dr. Joana Delgado Martins, Dr. José María Mateos Melero and Johannes Riemann from the Center for Microscopy and Image Analysis (ZMB) at the University of Zurich UZH for their kind help with image acquisition and analysis. We are grateful to Dr. Erik Slabber for his technical advice on cloning and PLA and help during manuscript preparation, and to our fellow Polymenidou lab members for the continuous and fruitful discussions on the project.

## Author contributions

M.P.-B performed plasmid cloning unless stated otherwise below, developed all stable isogenic HEK cell lines and carried out the corresponding experiments on them, including sample preparation for transcriptomics and proteomics. V.I.W. cultured the human neural networks, cloned LV transfer plasmids, produced lentiviral vectors and performed experiments on neurons and mutNLS isogenic HEK lines. A.Z. performed the full-length TDP-43-MBP protein purification, its corresponding plasmid cloning and *in vitro* experiments. L.D.V. provided cell culture support and performed immunocytochemistry experiments on isogenic HEK lines. C.F. performed triFC experiments on HeLa cells and its corresponding plasmid cloning. U.W. performed the RNA-seq analysis. I.M. and K.M.B. supported the RNA-seq analysis. A.C. performed the purification of TDP-43 RRMs and the NMR experiments. J.W. and Z.G. provided cell culture support. P.D.R. helped with the colocalization studies of nuclear markers and E.T. with the PLA. R.M. supported plasmid cloning and R.R. protein purification and *in vitro* experiments. S.S. and O.S. provided technical feedback. M.H.-P. provided samples of human neural networks, overall cell culture and experimental support and critical input on the study. F.H.-T.A., P.P. and M.P. provided supervision. M.P.-B, V.I.M., A.Z. and M.P. wrote and edited the manuscript and prepared the figures. M.P. directed the study. All authors read, edited and approved the final manuscript.

## Data and materials availability

All data needed to evaluate the conclusions in the paper are present in the paper and/or the Supplementary Materials. Proteomics data will be deposited to the ProteomeXchange Consortium via the PRIDE partner repository and the transcriptomics data will be deposited to the Gene Expression Omnibus (GEO) genomics data repository.

